# Chromosome level assembly of wild spinach provides insights into the divergence of homo- and heteromorphic plant sex-chromosomes

**DOI:** 10.1101/2023.07.17.549201

**Authors:** Edouard I. Severing, Edwin van der Werf, Martijn P.W. van Kaauwen, Linda Kodde, Chris Kik, Rob van Treuren, Richard G.F. Visser, Richard Finkers, Yuling Bai

## Abstract

**Background:** Cultivated spinach (*Spinacia oleracea)* is a highly nutritional crop species of great economical value that belongs to a genus of dioecious plant species with both homomorphic and heteromorphic sex chromosomes. The wild spinach species *Spinacia turkestanica* and *Spinacia tetrandra* are important genetic sources for improving cultivated spinach and excellent material for studying sex chromosome evolution in plants. However, until now there were no publicly available genome assemblies for these species.

**Results:** Here we sequenced and assembled the genomes of *S. turkestanica* and *S. tetrandra* and performed a tri-way comparative analysis with *S. oleracea*. We show that many abiotic- and biotic stress related gene clusters have expanded through tandem duplication in *S. tetrandra* after it diverged from the *S. turkestanica* - *S. oleracea* lineage. Focussing on the sex chromosomes we found that the previously identified inversion distinguishing the *S. oleracea* male- and female-SEX DETERMINING REGIONs (SDRs) is conserved in *S. turkestanica*. Although, the SDRs of these two species coincides with the PSEUDO AUTOSOMAL REGION of *S. tetrandra* the gene content is only partially conserved and the genetic factors determining sex in these species might differ. Finally, we show that recombination suppression between the *S. tetrandra* X- and Y-chromosomes resulted in a highly degenerated Y-chromosome and started before the species diverged from *S. turkestanica* and *S. oleracea*.

**Conclusions:** We expect that the novel wild spinach species genomes are of great value to the breeding community and evolutionary biologist especially focussing on the evolution of sex chromosomes in plants.

## Background

In dioecious plant species individuals either produce male or female flowers. In some dioecious plant species sex is determined by environmental instead of genetic factors [1]. In other species sex is determined by a pair of homomorphic- or heteromorphic sex chromosomes.

Homomorphic sex chromosomes are karyotypically indistinguishable and are highly similar in both size and gene content [2]. In contrast, heteromorphic sex chromosomes can be distinguished karyotypically and differ vastly in size and gene content [2]. In XY systems, gradual recombination suppression between the X and Y chromosomes eventually leads to degradation of the Y chromosome [3]. Different models have been proposed that describe the evolution including the degeneration of the Y chromosome (e.g. [4, 5].

The stepwise recombination suppression is typically characterized by the presence of evolutionary strata which correspond to homologous X and Y regions that have diverged at different time points in evolution following recombination suppression [6]. It has been suggested that structural variations, in particular inversions play a pivotal role in locally suppressing recombination [7]. Indeed, structural variations have been linked to evolutionary strata in both animal and plant species [8, 9].

The degradation process of the Y-chromosome leaves detectable signals including sequence loss, accumulation of repetitive elements and accumulation of deleterious mutations [3]. For only a handful of plants species, studies have been performed that specifically interrogate these signals for understanding Y-chromosome evolution (e.g. [9–11]. The advancement of long read sequencing technologies has enabled the publication of several phased sex determining regions or entire homomorphic plant sex chromosomes (e.g. [12–14]. However, to the best of our knowledge *Simmondsia chinensis* is the only plant species to date with publicly available phased heteromorphic sex chromosomes sequences [15].

Cultivated spinach (*Spinacia oleracea)* is a crop species of high nutritional and economical value. The two close wild relatives of *S. oleracea, S. tetrandra* and *S. turkestanica,* are agriculturally important as they serve as sources for novel traits, especially regarding resistance to downy mildew caused by the oomycete *Peronospora effusa* (Grev.) Rabenh [16]. Spinach is also a species of great interest for studying plant sex chromosome evolution as the genus *Spinacia* contains both species with homomorphic- and heteromorphic sex chromosomes. It has been demonstrated that the homomorphic and heteromorphic pairs are derived from a common ancestral set [17]. Several studies focussing on the sex chromosomes of spinach have been published in recent years (e.g. [18]. However, until now, only full-length sex chromosome sequences of the cultivated spinach *S. oleracea* are available but not of its wild relatives *S. turkestanica* and *S. tetrandra*.

Here nanopore (ONT) and Omni-C sequencing technologies were used for assembling the genomes of *S. oleracea* (cv. Viroflay) and its two wild relatives *S. turkestanica* and *S. tetrandra*. We performed tri-way comparative analysis between the cultivated and wild spinach species and specifically focused on two specific topics.

First, we briefly analyze the contribution of tandem duplication events in the creation and expansion of gene clusters in the spinach lineages as a whole and the individual spinach species.

Second, we performed an in-depth comparative analysis of the spinach sex chromosomes. We started by investigating the macro structure evolution of the homo- and hetero morphic sex chromosomes and searched for ancestral genes in the SEX DETERMING REGION (SDR). We then focused on determining when the heteromorphic sex chromosomes of *S. tetrandra* started to evolve and how the Y-chromosome is degenerating.

## Results

### Genome assembly

The highly repetitive genomes (79,9% to 84,1%) of a female cultivated spinach *S. oleracea* var. Viroflay and two male individuals of its wild relatives *S. turkestanica* and *S. tetrandra* were sequenced using Oxford Nanopore Technology (ONT) and subsequently scaffolded with Dovetail Omni-C illumina reads into six pseudo chromosomes for *S. oleracea* and *S. turkestanica* and seven for S*. tetrandra*. During course of this study, a chromosome level assembly of a female S*. oleracea* Viroflay was published [19]. We have decided to proceed with our assembly in this study for consistency in the assembly- and annotation methods. Furthermore, we followed the chromosome naming of the first released *S. oleracea* SPOv3 draft genome [20].

The assembled genome sizes (representing ∼89% to ∼95% of kmer-based estimated sizes) of *S. oleracea* and *S. turkestanica* are quite similar when compared to that of *S. tetrandra* (Additional file 1: Table S1). The much larger genome assembly of the latter species can mostly be attributed to the additional Y-chromosome (338,5 Mb pseudo chromosome and 24 Mb in additional related scaffolds).

The initial assembly size of the *S. tetrandra* Y chromosome was ∼243 Mb. However, mapping of illumina reads of the same male *S. tetrandra* individual revealed several regions on the X chromosome that had twice as high the expected coverage (Additional file 1: Fig. S1) while the Y chromosome had the expected haplotype coverage. The largest region is around 65Mb and located at the top of the X chromosome. On the other hand, normal diploid coverage was observed along the X chromosome and no coverage along the Y when examining the reads of a female *S. tetrandra* sample (Additional file 1: Fig. S2).

The region with elevated coverage could potentially be a PSEUDO AUTOSOMAL REGION (PAR) where recombination continues between the X- and Y-chromosomes [21]. As a result, the X and Y chromosomes could be so similar in this region that they collapsed in the initial assembly explaining the elevated read coverage observed in the male sample. Using phased SNP data, we reconstructed the X and Y haplotypes in this region (see material and methods). As the PAR is expected to evolve differently than the rest of the Y-chromosome it was treated separately or excluded in this study where appropriate.

BUSCO analyses indicated acceptable completeness of the gene-structure annotation of all three genomes (≥95%) (Additional file 1: Table S1). As expected, due to its larger size, the *S. tetrandra* genome has much more predicted genes (8466 (∼26%) and 9519 (∼30%)) than both *S. turkestanica* and *S. oleracea*. The total gene size in the *S. tetrandra* genome is 18.8 % and 19.6 % larger than that from *S. oleracea* and *S. turkestanica*, respectively. However, the biggest contributor to the larger genome size of *S. tetrandra* are repeat elements (Additional file 1: Fig. S3).

We clustered the genes of the three species using OrthoFinder and determined the number of genes that are unique to each genome. The results indicate that around 7% of the *S. oleracea* and 9% of the *S. turkestanica* genes are unique. However, in *S. tetrandra* up to 20.5% of the genes are unique (Additional file 1: Table S1).

We investigated whether the relatively high number of unique genes in *S. tetrandra* can be attributed to the presence of the heteromorphic Y-chromosome (excluding the PAR). To this end, we compared the fraction of non-PAR Y-chromosome genes in the *S. tetrandra* genome to the fraction non-Y genes that are unique to this species. We repeated the analysis for all autosomes and the X chromosome. For the autosomes and X chromosome, no significant differences (hypergeometric test) were found between the fraction of genes in the complete set and the fraction of genes in the unique gene set. In contrast, the fraction of Y genes in the unique gene set is ∼1.4 times higher than in the complete gene set. This result is highly significant according to a hypergeometric test (p=1.1e-63).

Further analysis showed that the Y-linked unique *S. tetrandra* proteins are significantly shorter (Kruskal Wallis: P: 0; followed by dunn’s test: P < 0.05) than non-unique genes (Figure 1a) but not different from unique genes on other chromosomes. The four most enriched biological process GO terms in the of unique Y genes are related to transposition and DNA-synthesis (Figure 1b). A full list of enriched GO terms is provided in supplementary table S2 (Additional file 1).

**Figure 1.**
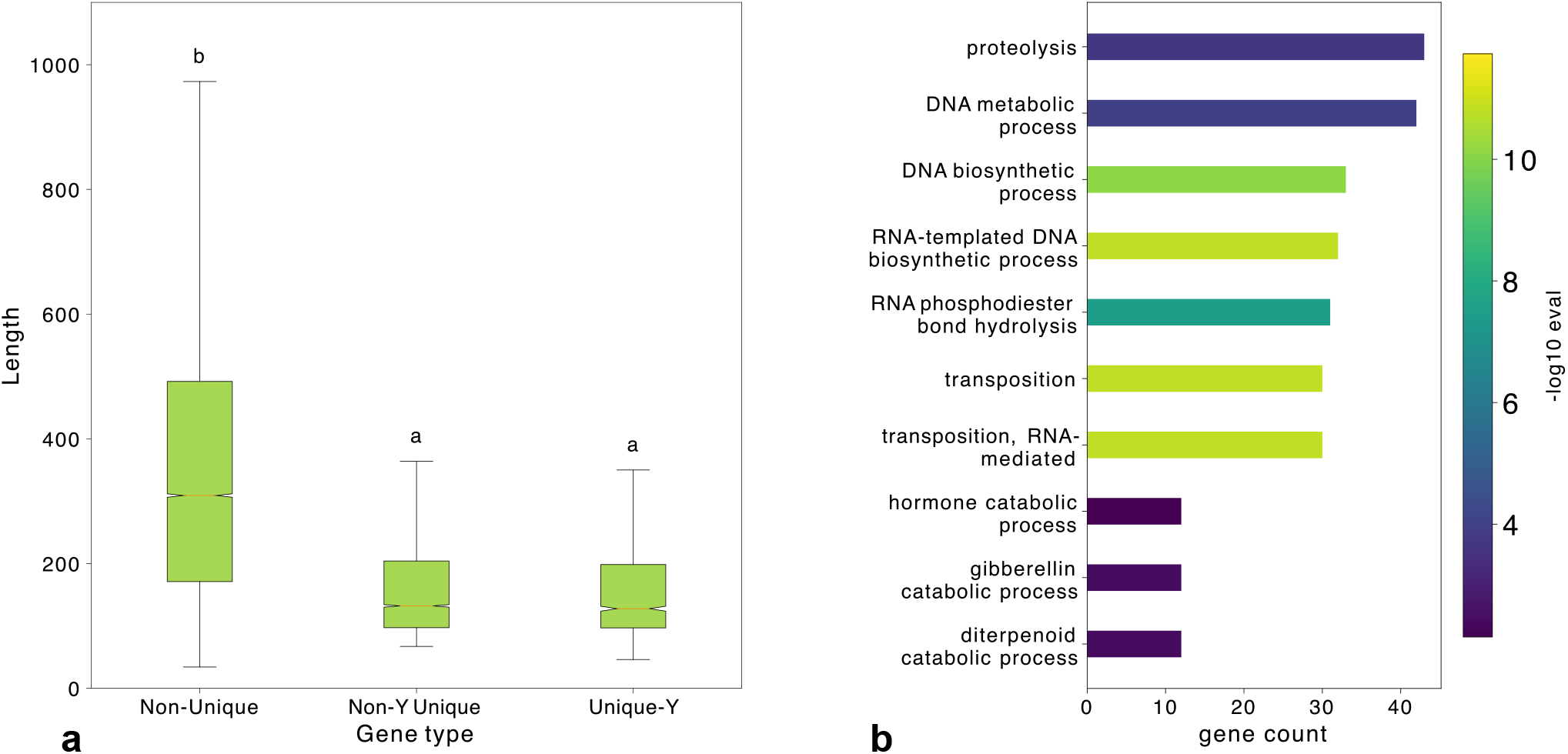
*S. tetrandra* protein lengths. **a** Length distribution for *S. tetrandra* proteins that have a homolog in at least one other spinach species (Non-unique), *S. tetrandra* specific proteins encoded on the Y-(Unique-Y) and other chromosomes (Non-Y Unique). Note, the total protein set analysed only contained the longest encoded protein sequence for each gene. Following a dunn post hoc test (P<0.05) the protein sets were divided into two groups (a and b). **b** Top 10 strongest enriched GO terms in the Unique-Y set.

### Tandem duplications

A major mode of adaption of plants to abiotic and biotic stress is through tandem duplications [22]. Especially recent tandem duplications in the wild spinach species can be of value for breeding purposes as they may comprise a source for novel traits such as resistance to pathogens. We inspected the contribution of tandem duplications to gene family expansion in each spinach species separately.

The number of genes in duplicated clusters we found using our method involved 6437 genes (2620 clusters) in *S. tetrandra*, 4833 genes (1921 clusters) in *S. turkestanica* and 5260 genes (2102 clusters) in *S. oleracea*. We performed GO terms enrichment analysis to determine whether a functional bias exists in the set of genes involved in tandem duplications. The analysis was performed separately in each species and a consensus list of GO terms enriched in all of them was made. As the result contained large numbers of similar GO terms we clustered the terms based on semantic similarity (see Methods). The largest cluster of related GO terms clearly indicates that genes involved in the response to biotic stimuli have been expanded through tandem duplications in the evolution of *Spinacia* (Figure 2).

**Figure 2.**
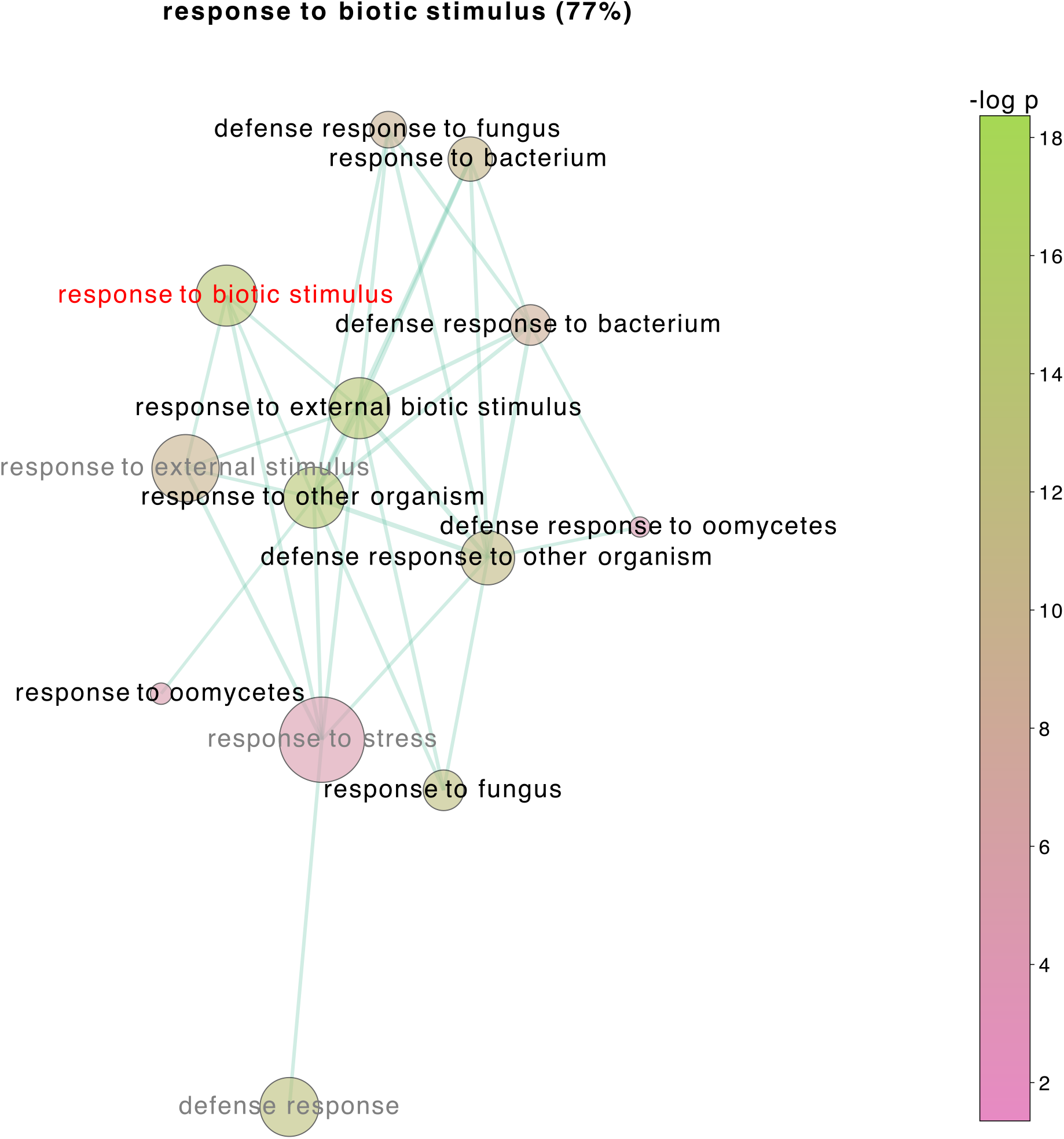
Enriched GO terms in tandem duplicated genes. GO-terms that were enriched (P < 0.01) in the tandem duplicated gene sets of all three species were clustered using pairwise semantic similarity (≥ 0.6). The largest cluster, shown here, is best represented by the term *response to biotic stimulus* (red). This term is a parent to 77% (black) of all cluster members. The remaining terms in grey do not descent from the representative term. Node size correspond to the number of tandem duplicated genes annotated to the term. Node colour reflects the strength of enrichment (−log10 P).

For the more recent tandem duplication events we used the gene duplication results from OrthoFinder as input for the detection of tandem duplication clusters. In total, 1522 *S. tetrandra* genes were found belonging to 701 clusters that were created or expanded through recent tandem duplication events. GO term analysis reveals that *S. tetrandra* has increased its repertoire of fungal infection response genes after splitting from the *S. turkestanica*-*S. oleracea* lineage. The most striking result is that out of the 67 enriched GO terms, 28 (42%) are transporters of varying bio molecules (Additional file 1: Table S3).

In total, 627 and 717 genes were found in 283 *S. oleracea* and 292 *S. turkestanica* clusters, respectively. However, unlike for *S. tetrandra* no enriched GO terms were identified for genes involved in recent tandem duplications.

### Genome structure comparison

Genome structure comparison between the three genomes (Figure 3a) demonstrates that *S. oleracea* and *S. turkestanica* only have limited structural differences when compared to *S. tetrandra*. On the other hand, *S. tetrandra* and *S. turkestanica* differ by several larger structural variations including inversions and a large reciprocal translocation between chromosome 2 and 6. It can also be observed that the *S. tetrandra* X- and Y chromosomes likely derived from a common ancestor shared with chromosome 1 of *S. oleracea* and *S. turkestanica*.

**Figure 3.**
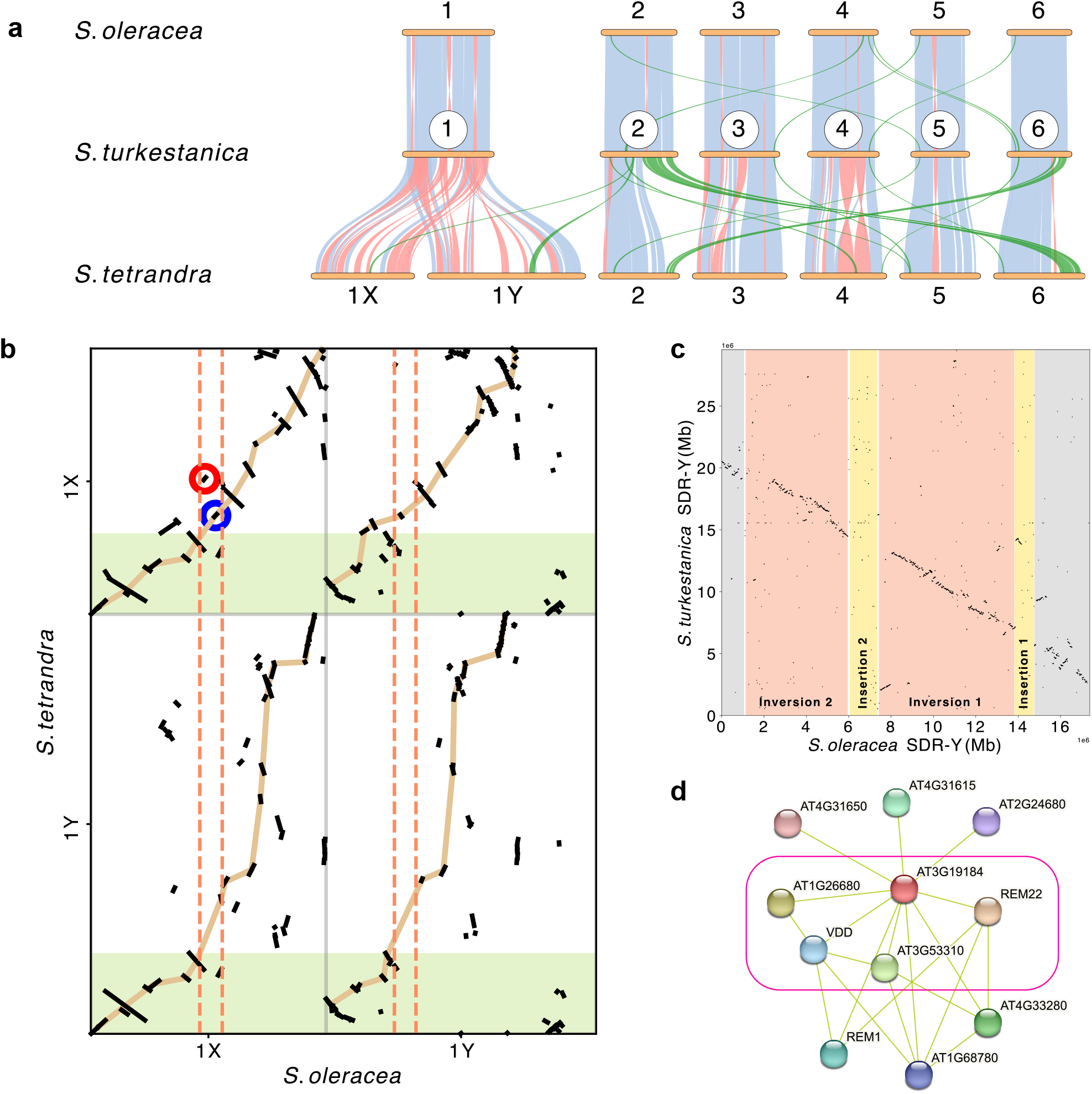
Genome structure comparison. **a** Macro structural comparison between the three spinach genomes. Co-linear blocks are connected by light blue lines. Pink and green lines indicate inversions and inter chromosomal translocations, respectively. **b** Structure comparison between the sex *S. tetrandra* and *S. oleracea* sex chromosomes. Beige lines represent the highest scoring monotonously increasing path though the syntenic blocks (black segments) as determined by dynamic programming. The PSEUDO AUTOSOMAL REGIONS on the *S. tetrandra* X- and Y-chromosomes are marked by a green rectangles. The boundaries of the *S. oleracea* X- and Y sex defining regions (SDRs) are delineated by dashed vertical orange lines. Red and Blue circles corresponded to specific conserved fragments referred to in the text. **c** Structural comparison between the *S. turkestanica* and *S. oleracea* Y-SDRs. Shaded regions correspond to structural variations between the X- and Y-*S. oleracea* SDRs as previously identified by Ma et al. In brief, regions on the *S. oleracea* Y-SDR that are co-linear with the X-SDR are represented in grey. Pink and yellow regions are inverted on and missing from the X-SDR, respectively. **d** Gene neighbourhood around the *Arabidopsis thaliana AT3G19184* gene as extracted from the STRING database. The neighbourhood is based upon co-occurrence of genes in PUBMED abstracts. A confidence score threshold of 0.5 was used for connecting nodes in the network. A subnetwork was created by single linkage clustering of nodes connected by edges representing scores of ≥ 0.7 (pink rectangle).

Direct comparison shows that besides the PAR, only at top of the *S. tetrandra* sex chromosomes there is a large co-linear block (Mb) (Additional file: Fig. S4) The remaining parts of the chromosomes are highly re-arranged. It can be expected that both the *S. tetrandra* X- and Y-chromosomes evolved separately in sequence and structure over the region where recombination seized. Therefore, we used the X-chromosome of *S. oleracea* as a reference point (Figure 3b).

After removing redundant alignments around 66,4% and 87% of *S .oleracea* X chromosome could be mapped to the *S. tetrandra* Y and X-chromosome, respectively (Additional file 1: Fig. S5). The other way around, around 68% of the *S. tetrandra* Y- and 82% of the S. tetrandra X chromosome could be mapped to the X chromosome of S. oleracea. (Additional file 1: Fig. S6). It was also observed that similar fractions of the *S. tetrandra* Y chromosome are translocated (27%) and inverted (22%) relative to the *S. oleracea* X-chromosome (Additional file 1: Fig. S6). On the other hand, almost half of *S. tetrandra* X-chromosome has been inverted in several individual events (Additional file 1: Fig. S6; Figure 3b).

Finally, of the conserved *S .tetrandra* Y and *S. olereacea* X fragment pairs, the *S. tetrandra* segment was more often than expected larger (Additional file 1: Fig. S7). No such size bias was observed when comparing the X-chromosome segment pairs (Additional file 1: Fig. S8).

### Conserved structural variation in sex determining regions of S. turkestanica and S. oleracea

Recently, it was shown that the homomorphic sex chromosomes of *S. oleracea* differ by a pair of large inversions and male specific insertions in the Sex Determining Region (SDR). The inversions was estimated to have occurred at around 1.9 Million years ago (MYA) [13]. Based on the synonymous substitution rate (K_s_) distribution of 4014 S. *olerecea* and *S. turkestanica* orthologous gene pairs we estimated that the two species have diverged around 0.2 MYA years ago (Additional file 1: Fig. S9). Even if the divergence age is likely an overestimate given that K_s_ values of zero were not considered, the species are predicted to be younger than the inversion event. We therefore investigated whether the sex specific structural variation found in *S. oleracea* is also conserved in *S. turkestanica*.

Given that the male *S. sturkestanica* individual that was sequenced in this study has an X Y chromosome pair we expected difficulties in assembling the Y SDR and therefore followed a non-trivial assembly approach (see Materials and Methods). The refined assembly of the SDR region resulted in contigs that when compared to the female *S. oleracea* SDR appear to have captured the breakpoints of an inversion (Additional file 1: Fig. S10). We therefore conclude that the inversion event is indeed ancestral to *S. oleracea* and *S. turkestanica* and has been conserved. Despite a number of smaller structural variations, the assembled SDR region is mostly co-linear to the *S. oleracea* Y SDR (Figure 3c).

Our SDR assembly does not contain the male specific insertions of *S. oleracea* Y-SDR (Figure 3d). In order to determine whether this a consequence of the assembly procedure or those sequences are specific to *S. oleracea* we analysed the read coverage of *S. turkestanica* ONT reads along the Y-SDR from *S. oleracea* (Additional file 1: Fig. S11). The analysis revealed that region corresponding to the insertions have half the expected diploid coverage and are not zero. We therefore conclude that these Y-specific insertion sequences are also present in the male *S. turkestanica*.

### Origin of the S. turkestanica and S. oleracea SDR

Given that the structural variation that distinguishes the male and female SDRs of *S. oleracea* and *S. turkestanica* preceded their divergence we wondered whether the SDR region is conserved in *S. tetrandra* and perhaps even ancestral to the spinach species. To this end we performed a synteny analysis between the sex chromosomes of *S. oleracea and S. tetrandra*.

The synteny analysis shows that the boundaries of the *S. oleracea* SDR correspond to regions located at the PAR boundary in *S. tetrandra* (Figure 3b). However, the central part of the SDR appears mostly absent from both sex chromosomes of *S. tetrandra*. The absence of this region was confirmed in a XX *S. tetrandra* assembly and is therefore unlikely the result of an assembly error due to XY chromosome-pairs within the PAR region (unpublished data).

Interestingly, two small syntenic regions of six (Additional file 2: Table S4) and ten (Additional file 2: Table S5) matched gene pairs were detected that reside within the *S. oleracea* SDR but are transposed in the *S. tetrandra* X-chromosome (blue and red circles in Figure 3, respectively).

Based on the eggnog function assignment and BLAST searches against the *Arabidopsis thaliana* Araport 11 database, one gene (*Stet|g4018.t1*), a B3 domain containing protein (Additional file 2: Table S4) was identified in the first block. The two best BLAST hits of this gene in the *A. thaliana* genome differ by only 7 bits in the score (183 and 176) while the score of the third best hit was 71 bits lower. We therefore considered to follow up on the two best hits – *AT5G42700* and *AT3G19184* which are both annotated as *AP2/B3-like transcription factor family protein*. The functional description in the STRING database [23] of these two genes indicate that they are also expressed in flower organs but is still rather broad (see Additional file 1: Appendix S1 and S2). In order to get more insight in the possible role of these genes we explored in which biological processes they are potentially involved by inspecting their association with other genes with known functions in the STRING database.

*AT3G19184 is located* within a network of 11 nodes which are linked because they are co-mentioned in multiple PubMed [24] abstracts (Figure 3d). By performing single linkage clustering of the nodes using only the high confident edges (>0.7) *AT3G19184* linked 4 other genes (Pink rectangle in Figure 3d). Three of these genes are of particular interest: *REM22* is expressed in stamen primordia and the placental region of developing carpels and the ovary. *AT1G26680* is expressed in the ovule and finally *VDD* is a direct target of the MADS domain ovule identity complex. Altogether this high confident sub network suggests that *AT3G19184* and perhaps *S. tetrandra Stet|g4018.t1* function in reproductive organ formation. Furthermore, the ortholog of this gene in *S. oleracea* was found to have a sex dependent expression pattern in developing flowers [13] (see additional file 1: Figure S12a). More specifically, while the expression levels are not significantly different in the initial stages of developing flowers, the expression level strongly drops in male flowers but not female flowers at stage 3 where ovary starts to differentiate [13].

The network around *AT5G42700* only shares *AT1G26680* with the network around *AT3G19184* (Additional file 1: Figure S13). The function of genes in this network are less specific (Appendix S2).

Further syntheny analysis shows that the order of genes around *AP2/B3* like is disrupted on the *S. tetrandra* Y chromosome but four of the genes (including the gene of interest) are in conserved order on chromosome 1 of *B. vulgaris* (Additional file 2: Table S4). The latter indicates that the gene was already present in the ancestor of spinach. The protein sequence encoded by the *B3* domain containing genes is highly conserved (Additional file 3: Fig. S14). The only noteworthy difference within the spinach species is the 3 amino acid gap and different C terminus of the protein encoded on the Y chromosome of *S. oleracea*.

The second syntenic block (Additional file 2: Table S5) is mostly colinear between *S. oleracea* and chromosome X of *S. tetrandra* (10 genes). Only small pieces can be considered co-linear on the *S. tetrandra* Y-chromosome (5 genes) and even less on *B. vulgaris* (3 genes).

*S. tetrandra Stet|g3474.t1*) is one of the three genes in conserved synteny and annotated. The encoded protein is annotated as a *Plant protein of unknown function (DUF868)* and has a clear single best BLAST hit *AT2G04220* in *A. thaliana*. Unfortunately, to date no neighbourhood information exists in the STRING database for this gene. However, according to the TAIR10 annotation this gene is expressed in pollen and pollen tube cells. This gene is also well conserved, but in contrast to the *B3* domain containing gene, has no obvious differences in the spinach species (Additional file 4: Fig. S15). However, the gene is activated between stage 4 and 5 of only male flower development in *S. oleracea* (additional file 1: Fig S12b). At stage 4 distinct anthers are form in male *S. oleracea* flowers [13].

Ma and colleagues [13] proposed two candidate genes (NRT1/ PTR FAMILY 6.4 and eukaryotic translation initiation factor 3 subunit A-like) for male determinacy within a genomic region specific to the Y-SDR of *S. oleracea* containing 17 genes. In order to determine when this Y-specific region originated we searched the proteins encoded by the genes within this region in our genome assemblies including that of *B. vulgaris* using exonerate.

The candidate genes were only found intact and adjacent to each other on chromosome 0 (un-anchored contigs) of *S. turkestanica* but no good matches were found in our female *S. oleracea*, *S. tetrandra* and *Beta vulgaris*. Several of the genes in Y-specific region upstream of the candidate genes are also found in a block (albeit a bit re-arranged) in our female *S. oleracea* (Additional file 2: Table S6) and *S. turkestanica* (Additional file 2: Table S7). On both sex-chromosomes of *S. tetrandra*, only three of the upstream genes are found close to each other and in *B. vulgaris* just 2 on chromosome 4 (Additional file 2: tables S8-S10). No colinear block was found corresponding to the Y-specific region downstream of the candidate genes.

### Age of the S. tetrandra heteromorphic sex chromosomes and evolutionary strata

In order to determine the age of the *S. tetrandra* sex chromosomes we first identified 763 ortholog groups that consisted of exactly one *S. oleracea* and *S. turkestanica* gene linked to chromosome 1 and exactly one non-PAR X and Y linked copy in *S. tetrandra*. We generated a distribution from pairwise K_s_ values between the *S. tetrandra* X and Y genes in 756 of these orthogroups.

We modelled the distribution of K_s_ values as a mixture of *n* Gaussian components with *n* being based upon Bayesian Information Criterion (BIC) after fitting multiple Gaussian mixture models with varying number of components (see material and methods). The modes of the individual Gaussian components were used for calculating divergence times between X and Y gene pairs (see material and methods). The same procedure was repeated using 11237 *S. tetrandra*-*S. oleracea* autosomal ortholog pairs for getting an estimate of the divergence time between the two species. Based on the BIC, two components were selected for both the *S. tetrandra*-*S. oleracea* (Additional file 1. Fig. S16) and the *S. tetrandra* sex chromosome – linked (Additional file 1: Fig. S17) gene sets. Given the modes of fitted Gaussian distributions on the ortholog K_s_ data, we estimate that *S. tetrandra*-*S. oleracea* diverged around 5.2 MYA ago (Figure 4a). The second height of the second fitted distribution was so small that it was discarded (Additional file 1: Fig. S18). For the sex-linked genes of *S. tetrandra* we the modes of the two fitted Gaussian distributions translated to 3.3 and 8.0 MYA (Figure 4a; Additional file 1: Fig. S19). These results suggest that different regions of the *S. tetrandra* sex chromosomes started to diverge at different time points.

**Figure 4.**
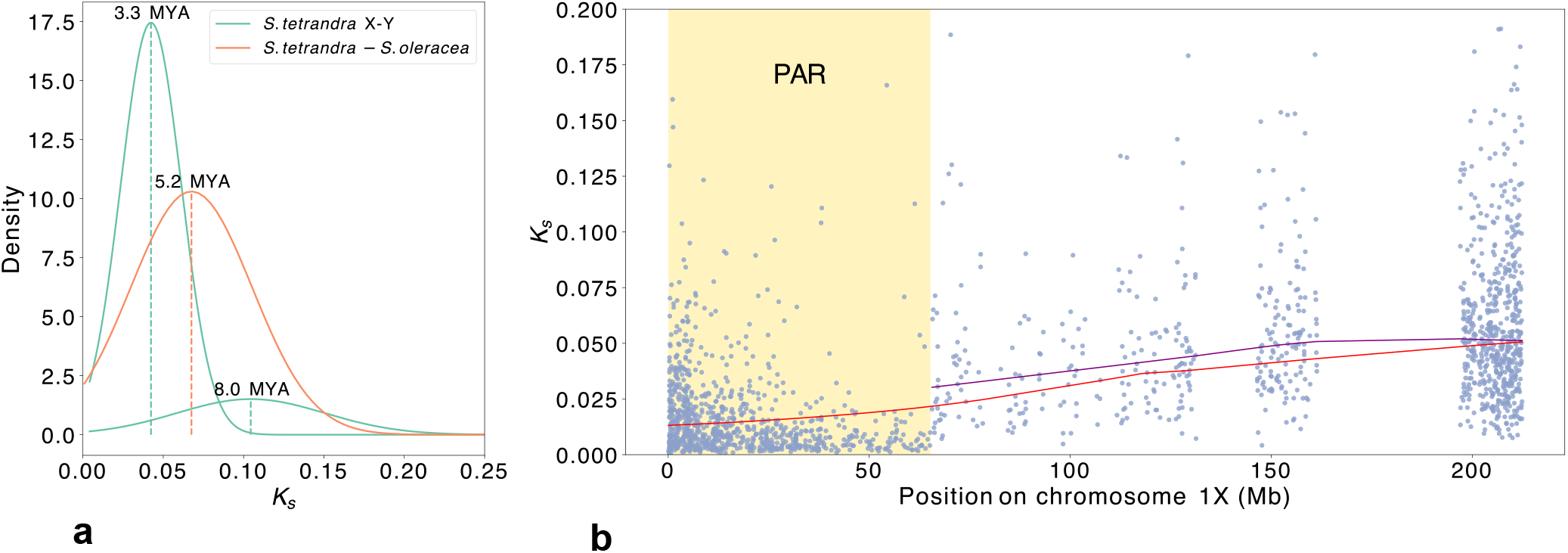
*S. tetrandra* sex chromosome divergence. **a** Individual Gaussian mixture components (distributions) underlying the empirical distributions of synonymous substitution rates (K_s_) for *S. tetrandra* X-Y homolog- and *S. tetrandra*-*S. turkestanica* autosomal ortholog pairs. The K_s_ values at the modes of the individual components were used for calculating divergence times in Million years ago (MYA). **b** Pairwise K_s_ values for *S. tetrandra* X-Y homolog pairs plotted along the X-chromosome. The PSEUDO AUTOSOMAL REGION (PAR) is highlighted in yellow. The red trendline across the chromosome was obtained by performing LOWESS regression on all K_s_ values. A second trendline (purple) was generated using only the pairwise K_s_ values of non-PAR gene pairs.

Stepwise divergence of different blocks along sex chromosomes lead to the formation of evolutionary strata [6]. In order to determine whether this is also the case in *S. tetrandra* we plotted the K_s_ values of 1870 X-Y orthologs within conserved syntenic regions along the X chromosome. We indeed observed a tendency for K_s_ values to increase from the PAR towards the end of the chromosome (Figure 4b).

### S. tetrandra Y-chromosome size and LTR ages

Comparison of size and composition of *S. tetrandra* chromosomes shows the Y-chromosome is much larger than the rest mainly due to the higher amounts of LTR elements (Figure 5a). This could mean that either the Y chromosome is more prone to LTR insertions or that the Y chromosome less efficient at purging the LTR elements. Although the former is difficult to test, the latter can be tested under the following assumptions (I) each chromosome is equally likely to be targeted by a retrotransposon, (II) transposons are continuously spreading (including in bursts). (III) pruning of transposons by recombination occurs at similar rates on all normally recombining chromosomes. Under these simplified assumptions we expected that autosomes including the X chromosome are likely to have similar LTR element age distributions while the Y chromosome, excluding the PAR, would be (slightly) more biased towards older elements due to inefficiency of removing elements.

**Figure 5.**
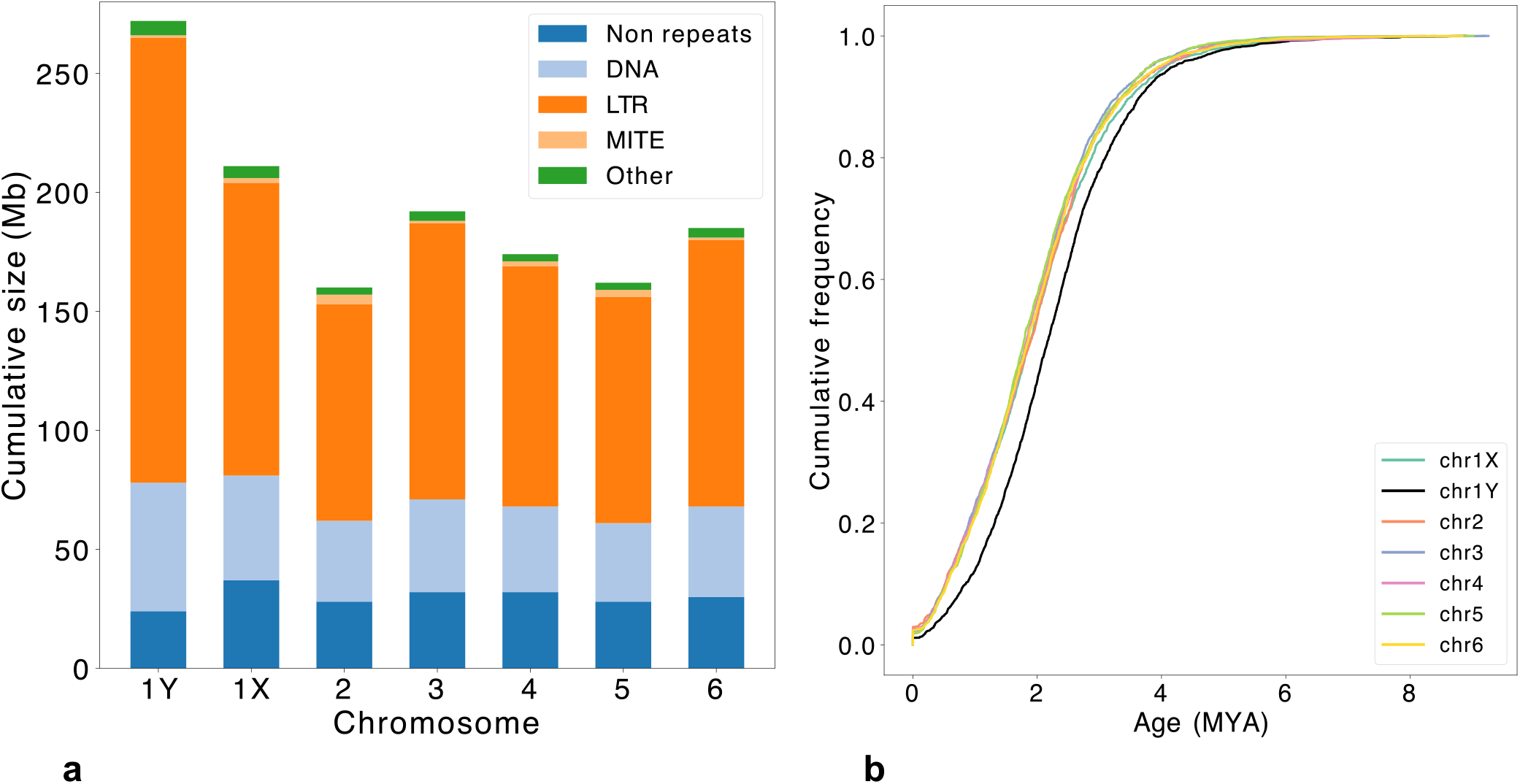
*S. tetrandra* repeats. **a** Repeat element composition per chromosome. Note that due to their very low summed sizes, LINE elements were incorporated into the “Other” repeat category. **b** Chromosome specific cumulative frequency distributions of LTR insertion ages. The distribution for the Y chromosome (black) is the only one that is visually separable from those of the other chromosomes.

According to pairwise Kolmogorov–Smirnov tests (Additional file 1: Table S11) the empirical LTR insertion age distributions of all autosomes significantly lie above that of the Y chromosome (Figure 5b) indicating a bias on the latter for older elements. Additional Kruskal Wallis test of the LTR ages also suggests a significant difference in LTR age groups (p <2.2e-16). Follow up Dunn tests (Additional file 1: Table S12) indicates that the chromosomes can be divided into two groups with similar median LTR ages (Additional file 1: Fig. S20). The first group contains all autosomes included the X chromosome and the second contains only the Y-chromosome.

### Accumulation of protein mutations on S. tetrandra sex chromosomes

We tested the hypothesis which suggests that reduced recombination of the Y-chromosome results in less efficient purifying selection and therefore increased accumulation of mutations relative to the X chromosome. To this end, we determined the number of amino acid changes of 763 *S. tetrandra* X- and Y-linked gene pairs relative to their *S. oleracea* ortholog (Figure 6)

**Figure 6.**
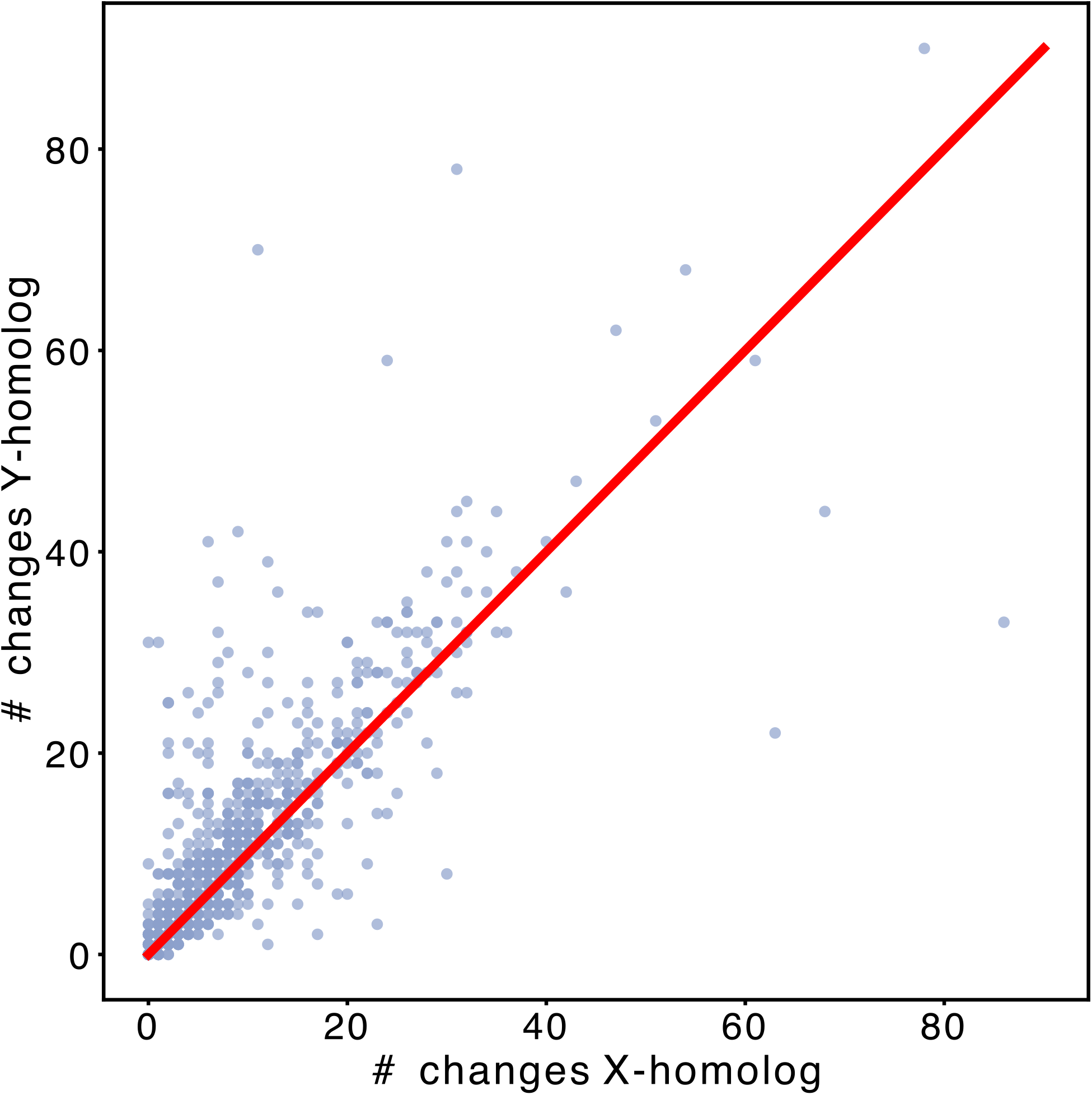
Amino acid changes of *S. tetrandra* X-Y protein pairs relative to *S. turkestanica*. The red line (*y=x*) corresponds to cases where the X- and Y-proteins have equal number of amino acid changes relative to *S. turkestanica*. Dots above the line represent protein pairs for which the Y-protein has more amino acid changes than its X-counterpart. The opposite is true for dots below the red line.

Out of the 323303 aligned amino acid residues positions (without gaps), 7393 (2.3%) and 9055 (2.8%) amino-acid changes were found for the Y and X-linked *S. tetrandra* proteins relative to the *S. oleracea* ortholog, respectively. The difference in protein mutation accumulation rate is small (1.2x), but significant (G-test, p <2.2e-16).

In order to ensure that the observed difference in amino acid changes is not just the result of several proteins having excessive number of amino acid changes, we quantified how often the *S. tetrandra* Y protein accumulated more mutations than the *S. tetrandra* X protein. In 462 (70%) of 660 (excluding ties) protein pairs, the number of accumulated mutations of *S. tetrandra* Y protein is higher than that of the X protein. This result is highly significant (p= 1.5e-25) according to a binomial test with success rate 0.5.

### Pseudo genes

Another effect of mutation accumulation is the potential to introduce premature stop codons or frameshifts into the open reading frame of genes leading to loss of function/pseudogenization. Given that Y-linked genes in *S. tetrandra* have accumulated more mutations than their X-linked counterpart we checked whether pseudogenization is more frequent on the Y-chromosome. To this end, we determined what fraction of X- and Y genes have corresponding pseudogene fragments with open reading frames that are disrupted by either frame shifts or premature stop codons. More specifically, for each X- and Y gene we searched for homologous gene fragments on the other chromosome that do not overlap with any annotated gene and that subsequently pairs with the query gene in a collinear gene block (see material and methods). We performed the analysis separately for genes within and outside the PAR. For the analysis with non-PAR X genes as queries a total of 101 gene matches were found of which 56 (55.4%) were disrupted by either a frame shift or internal stop codons. A similar fraction (54.7%; 52 out of 95) were found when using the non-PAR Y genes as queries. For PAR genes, 15.5% (26 / 168) and 15.1% (23 / 152) cases were disrupted for X- and Y queries, respectively.

## Discussion

In this study we assembled the genomes of one cultivated and two wild spinach species. Previous studies have shown that *S. oleracea* and its wild ancestor *S. turkestanica* are very closely related [17]. It was therefore expected that the assembled genomes of the two species are very similar in terms of size and gene content. *S. tetrandra* has a much larger assembled genome resulting mainly from the presence of the large additional heteromorphic Y chromosome. We estimate using re-sequencing data that the PSEUDO AUTOSOMAL REGION of *S. tetrandra* X- and Y chromosome where they still recombine is around 65Mb.

The *S. tetrandra* Y-chromosome was found to be enriched for unique genes that did not cluster with any *S. oleracea* or *S. turkestanica* genes. The unique genes are significantly much smaller than non-unique and could be species specific micro-proteins [25]. However, the fact that they are enriched for transposition and DNA-synthase functions can also indicate that many might represent transposon fragments that have escaped repeat masking and were subsequently annotated as genes.

Consistent with previous studies e.g. [22] we found gene families that have expanded through tandem duplications are significantly enriched for GO terms involved in the response to biotic stimuli. Only for *S. tetrandra* did we find significant GO term enrichments, including response to fungi and herbivores, for tandem duplicated genes that originated after divergence from *S. oleracea* and *S. turkestanica*. Of special interest are the large number of enriched terms relating to various molecular transporters which may play important roles including plant development and abiotic stress resistance [26]. These recent tandem duplicated gene clusters are of particular interest for breeding programs as they can serve as a genetic source for crop improvement. The fact that no functional enrichment was found for recent tandem duplicated genes of *S. turkestanica* and *S. oleracea* can be explained by their very recent divergence.

Macro synteny analysis between the three species confirms that heteromorphic *S. tetrandra* chromosomes are derived from a common ancestral chromosome pair [17]. Unlike, the *S. tetrandra* Y chromosome which appears highly fragmented, the X-chromosome has higher sequence similarity with the sex chromosomes of *S. oleracea*. Consistent with other plant species e.g. papaya [9] and the other two spinach species (see below) inversions are also implicated in the sex chromosome divergence in *S.tetrandra*.

The female and male SDRs of *S. oleracea* have previously been shown to also differ by large inversions that occurred around 1.9 MYA [13]. Despite assembly difficulties arising from sequence a XY individual we were able to show that these inversions occurred before the *S. oleracea* and *S. turkestanica* diverged.

Further back in evolution, the region adjacent to *S. oleracea* / *S. turkestanica* SDRs are homologous to the predicted *S. tetrandra* PAR. This suggests that the initial sex determining region probably were the same in *S. tetrandra* and the lineage leading to *S. turkestanica/S. oleracea*.

The two small syntenic fragments of genes conserved in the SDRs of *S. turkestanica*, *S. oleracea* and the sex chromosomes of *S. tetrandra* was already (partially) present in *B. vulgaris* and contain two genes of interest. One of those genes is predicted to be a *B3 domain* containing protein that resembles an *Arabidopsis* AP2/B3 protein which in turn is predicted to function in sex organ differentiation. While the male *S. oleracea* version of the protein has a 3 amino acid deletion, the male *S. tetrandra* version has no major changes compared to the female versions. Nevertheless, given the sex-biased expression levels of this gene in *S. oleracea* tentatively suggest that this gene might be important for the proper development of female flowers. The second candidate gene is one of unknown function but its best hit in Arabidopsis thaliana is expressed in the pollen and pollen tube cells. Indeed, the protein encoded by this gene is upregulated during anther maturation of male flowers in *S. oleracea*.

We found the candidate genes for male determination in *S. oleracea* proposed by Ma and colleagues [13] in *S. turkestanica* but not in our female *S. oleracea*. However, we did find that the region upstream of these candidate genes is also present in the female *S. oleracea* and thus not Y-specific. The lack of good synteny around and the absence of the candidate genes in *S. tetrandra* and *B. vulgaris* indicates that they are novel. Therefore, different regulators might determine male sex differentiation in *S. tetrandra* and the lineage leading to *S. turkestanica* and *S. oleracea*.

We used synonymous substitution rates between *S. tetrandra* X-linked and Y-linked orthologs for inferring the age of the sex-chromosomes. Our analysis indicates that the empirical K_s_ distribution can be modelled by a mixture of 2 Gaussian distributions with modes that translate to 8.5 and 3.3 MYA. This result suggests that the *S. tetrandra* X- and Y-chromosome did not diverge in a single event but in at least two episodes. Our estimate of the divergence time between *S. oleracea* and *S. tetrandra* is 5.2 MYA which is not too different from the 5.66 MYA reported previously [27]. Therefore, we conclude that parts of the *S. tetrandra* X- and Y-chromosomes started to diverge before the species split the common ancestor *S. oleracea and S. turkestanica*.

As recombination is gradually suppressed along the length of the Y-chromosome, different parts will start diverging from their homologous X chromosome regions at separate points in evolution leading to strata [28]. Evolutionary strata have previously been identified in animals (e.g. [8, 29] as well as in plant species (e.g. [9–11]. In agreement with previous studies, we also observed a gradient in gene divergence along the X chromosome.

Increase in size due to the accumulation of repetitive elements has been described as one of the consequences of recombination suppression [3]. Indeed, we observed that the total base count of repetitive elements and in particular LTR-elements on the *S. tetrandra* Y chromosome is almost as large as the size of each chromosome except the X chromosome. We also found that intact LTR elements on the Y chromosome tend to be biased towards older ages compared to those on other chromosomes. It has been shown that recombination rate in plants is negatively related to LTR content in genomes [30]. Therefore, the bias towards older elements potentially reflects that due to recombination suppression the Y chromosome has become less efficient in removing LTR-elements.

We have shown that the *S. tetrandra* genes on the Y chromosome have significantly accumulated more non-synonymous mutations than their X-chromosome homolog. These results support the hypothesis that the lack of recombination can lead to a decrease in selection efficacy [11, 31]. However, unlike in *Silene latifolia,* where Y genes were shown to more rapidly pseudogenize than X genes [10], we find no such difference. The rates only differed between genes within and outside the PAR.

## Conclusions

In this study we present the genome sequences of cultivated spinach and its two closely related wild species. We showed the potential of especially *S. tetrandra* as a genetic source for crop improvement. We also used the assemblies to investigate the complex evolution of sex chromosomes in the genus *Spinacia*. Our study was not designed to and cannot indicate which of the competing mechanistic models describing the processes of Y chromosome degeneration best applies to spinach. However, we did find the tell-tale signs that the *S. tetrandra* Y chromosome is degenerating. Finally, we believe that these genomes will be of great value to the plant sex research and spinach breeding communities.

## Methods

### DNA extraction and preparation

High molecular weight DNA for long-read sequencing was isolated from fresh young leaves of *S. oleracea* cv. ‘Viroflay’, and a male and female genotype from each of the wild relatives *S. tetrandra* MGK12 (CGN25471) and *S. turkestanica* KK28 (CGN25259), using a modified version of the Bernatzky and Tanksley protocol [32]. The two accessions from the wild spinach species were collected by the Centre for Genetic Resources, the Netherlands (CGN) during collecting missions in 2008 in Central Asia and in 2011 in the Trans Caucasus [33]. Accession information can be accessed on CGN’s website [34].

### Nanopore sequencing

Libraries were prepped using the 1D ligation sequencing kits SQK-LSK109 and SQK-LSK110 (Oxford Nanopore Technologies, Oxford, UK) according to standard protocol. The *S. turkestanica*, *S. tetrandra* and *S. oleracea* samples were sequenced on an Oxford Nanopore GridION using resp. 9, 7 and 4 flow cells using the standard protocol, yielding ∼90Gbases, ∼73Gbases and ∼37Gbases, respectively. Base calling was performed using Guppy v3.2 (Oxford Nanopore Technologies).

### Illumina sequencing

DNA of the spinach genotypes was used to generate 150-bp paired-end reads on an Illumina HiSeq 2500 (GenomeScan, Leiden, The Netherlands). Samples were processed using the NEBNext^®^ Ultra DNA library Prep Kit from Illumina. Genome characteristics were estimated using Jellyfish v2.2.10 [35] k-mer counts and GenomeScope [36].

### Nanopore reads pre-processing

Porechop [37] was used to remove the adapter sequences from the reads and Filtlong [38] was used to remove short reads . The remaining reads were mapped against the chloroplast and mitochondrial sequences of the corresponding species, and all reads with a mapping length > 45% of the read length were discarded.

### Organelle assembly

Circular chloroplast sequences of *S. tetrandra* and *S. turkestanic*a were assembled using the GetOrganelle pipeline [39] with a 4M subset of the Illumina reads using default settings.

In order to assemble the mitochondrial genome, we first mapped adapter trimmed nanopore reads larger than 50kb against a publicly available *S. oleracea* mitochondrial genome sequence (KY768855). Reads mapping over >70% of their length were identified using samsift [40] and extracted using seqtk [41]. The extracted reads were assembled using flye [42] with default parameters. The largest contig from the flye assembly was then used for a new iteration of mapping and extracting reads mapping >70% of their length. A subsequent assembly using flye resulted in a circular contig that was polished using two rounds of Pilon [43] with Illumina reads.

### Genome assembly

Processed Nanopore reads were assembled using the Nextdenovo program [44] (read_cutoff = 3k; read_type=ont). Nextpolish [45] was used to polish the contigs, using the long nanopore reads, and the Illumina reads from the same sample (relevant parameters: sgs_options = -max_depth 100 -bwa; lgs_options= -min_read_len 1k -max_depth 100; lgs_minimap2_options = -x map-ont) . Due to the heterozygosity of the accessions, some parts of the genome are assembled in separate contigs, containing both alleles. To identify and remove these redundant contigs we used purge_haplotypes [46] (default settings). After this step, most of the contigs contained both haplotypes. Finally, Haplo-G [47] was used to remove read-errors and to keep the reference sequence biased to one of the two haplotypes, to prevent haplotype switching of the sequence (default settings).

### Dovetail Omni-C Library Preparation and Sequencing

For each Dovetail Omni-C library, chromatin was fixed in place with formaldehyde in the nucleus and then extracted. Fixed chromatin was digested with DNAse I, chromatin ends were repaired and ligated to a biotinylated bridge adapter followed by proximity ligation of adapter containing ends. After proximity ligation, crosslinks were reversed and the DNA purified. Purified DNA was treated to remove biotin that was not internal to ligated fragments. Sequencing libraries were generated using NEBNext Ultra enzymes and Illumina-compatible adapters. Biotin-containing fragments were isolated using streptavidin beads before PCR enrichment of each library. The library was sequenced on an Illumina HiSeqX platform to produce an approximately 30x sequence coverage.

### Scaffolding with HiRise

The *denovo* assemblies and their corresponding Dovetail OmniC library reads were used as input data for scaffolding using HiRise [48]. Dovetail OmniC library sequences were aligned to the draft input assembly using BWA [49].The separations of Dovetail OmniC read pairs mapped within draft scaffolds were analyzed by HiRise to produce a likelihood model for genomic distance between read pairs, and the model was used to identify and break putative miss-joins, to score prospective joins, and make joins above a threshold.

### Ordering and renaming of the pseudo chromosomes

The final assembly scaffolds (pseudo-molecules) were mapped against the Spov3 genome [50] using minimap2 (−x asm20) [51]. The resulting alignments were used for orientating and numbering the chromosomes according to the Spov3 assembly. The sex-chromosomes are explicitly called 1X and 1Y in *S. tetrandra*.

### Refined phasing of S. tetrandra PSEUDO AUTOSOMAL REGION

Omni-C reads of the male sample were mapped against the initial X chromosome with BWA and phased using HapCUT2 [52]. Next, we used mapped ONT reads and illumina reads for extending the size of the initial HapCUT2 blocks using Whatshap [53]. The PSEUDO AUTOSOMAL REGION (PAR) was identified as genomic segments having 2x the expected read coverage after mapping illumina reads of the male *S. tetrandra* sample using BWA. The consensus sequence of the individual phases were extracted from the suspected PAR using bcftools [54]. The mapped reads from both the male and female *S. tetrandra* individuals were used for labelling phases as either X- or Y. The sequences from the X-phase were used for modifying the PAR on the X-chromosome.

The final Y-chromosome was generated by remapping Omni-C reads to the initial Y-chromosome including phased sequences labelled as Y. SALSA2 [55] was used for scaffolding the final Y-chromosome.

### Re-assembling the SEX DETERMINING REGION of S. turkestanica

Nanopore reads were mapped to the recently published YY *S. oleracea* genome [13] using minimap2 (−x asm5). All reads mapping to chromosome 1 were extracted and corrected using the read correction module of NECAT [56] . The corrected reads were subsequently assembled using NextDenovo. NextPolish was then used with the nanopore reads for cleaning the initial contigs. The polished contigs were mapped to both the initial *S. turkestanica* assembled chromosome 1 and the YY *S. oleracea* chromosome 1 using minimap2. The resulting alignments were fed into custom python scripts that produced an initial ordering of the contigs that was manually inspected and corrected.

### Quantifying expression levels of potential SEX related genes

RNA-seq time-series corresponding to developing male and female flowers performed by Ma and colleagues [13] were downloaded from NCBI Sequence Read Archive [57] (Project number: PRJNA724923). Gene expression levels were first quantified using Salmon [58]. Stage specific differential gene expression (adjusted p-value threshold 0.01) analyses were performed using DESeq2 [59].

### Repeat annotation

For each genome, a *de novo* transposable element (TE) library was created using EDTA [60] in sensitive mode. Intact LTRs were extracted from the EDTA output. The TE libraries produced by EDTA were subsequently used for masking the corresponding genome using RepeatMasker with the following parameter settings: -no_is -nolow -e rmblast. [61] .

### Age of LTR retrotransposons

For calculating the insertion age of intact LTR retrotransposons, the long terminal repeats of the retro transposons were aligned using muscle [62]. The dist.dna function of the ape R package [63] was used for calculating the Kimura 2 parameter distance [64] (K80 model) between the long terminal repeats. Conversion of the Kimura distance (K) to years was done using the formula K / 2r [65] where r corresponds to the number of substitutions per year which was set to 6.5e-9 [27]

### Gene annotation

BRAKER2 [66] was used in two different modes for de novo gene-structure annotation. For the protein mode of BRAKER we used OrthoDB v10 [67] plants proteins dataset as query. For the RNA-seq mode, we downloaded paired end RNA-seq reads (PRJNA663445) corresponding to different *Spinacia oleracea SP75* tissues from the NCBI Sequence Read Archive [57]. The reads were first mapped to the genome using STAR [68] and the resulting alignments were subsequently used as input for BRAKER2. The gene-structure predictions of the BRAKER runs were combined into a final annotation using TSEBRA [69]. Function- and GO-term annotations were assigned to the predicted gene models using eggNOG mapper [70].

### GO term enrichment analysis

GO term enrichment analysis were performed using the classic test of the TopGO R package [71]. Grouping of large number of closely related GO terms was done using the pairwise sematic similarity between GO terms as implemented in GO-Figure! [72]. GO terms with pairwise similarities of 0.5 or more were clustered using the mcl program [73] as it was found to give better results than single linkage clustering. Finally a representative term was selected for each cluster that was either: 1. a cluster member that is a parent term to 75% of the cluster members, or 2. the term of all possible parent terms to the cluster with lowest frequency in the Uniprot GOA database [74].

### Orthologous groups and syntheny analysis

Orthogroups were identified with OrthoFinder [75] using only the longest encoded protein for each gene and using the N0.tsv produced in the Phylogenetic_Hierarchical_Orthogroups folder.

All intra- and inter species synteny analyses were performed using MCScanX [76]. Three-way synteny plot was generated using custom scripts in the style of a karyotype plot from the JCVI utility library [77].

### Tandem duplications

Gene families within a species were constructed by first performing an all versus all BLAST [78] search (−e 1e-10, -F F). The blast hits were filtered by removing all HSPs involving less than 150 amino acids or with an alignment identity <30%. The remaining hits were used for clustering proteins using mcl (−I 2). Tandem duplicated proteins were considered as genes within a cluster that are separated by 4 genes at the most. For recent tandem duplications we used terminal gene duplications as identified by OrthoFinder.

### Synonymous substitution calculation

For calculating pairwise K_s_ values we first aligned the proteins using MAFFT [79] and removed lowly conserved regions using trimAL [80]. Low conserved regions were removed from the alignments using trimAL [80]. The resulting trimmed alignments were used as a guide to generate codon alignments using the original CDS sequences from the protein pair. After removal of gapped positions and stop codons, K_s_ values were calculated using the method of Nei and Gojobori [81] implemented in KaksCalculator version 2.0 [82].

### Calculating divergence times

In order to calculate divergence times between *S. oleracea* and *S. tetrandra* and between the sex chromosomes of *S. tetandra* we first created K_s_ distributions. Upon visual inspection we assumed that these distributions could be modelled using a single or mixture of multiple Gaussian distributions. The number of components for each distributions was determined by making mixture models with different number of components using the GaussianMixture class from scikit-learn python package [83]. The final number components were selected based upon Bayesian information criteria (BIC) values. In case no clear lowest BIC value was observed we used the elbow approach. During the course of the experiment, after multiple times running the modelling procedure, we occasionally obtained slightly different results that were considered the result of different starting points of the pseudo random generator. We therefore fixed the seed of the pseudo random generator of numpy prior to 12345 before each modelling procedure to ensure reproducibility of our results.

For the divergence time between *S. oleracea* and *S. turkestanica* we fitted a line using the Kerneldenstiy class of scikit-learn [83]. The best bandwidth was selected by exploring several bandwidths and selecting best one using the GridSearchCV class.

After fitting Gaussian models or Kernel density we searched for peaks using the find_peaks function from the scipy python package [84]. The K_s_ value corresponding to the peaks was translated to years using the same formula as for the LTR insertion time estimates. However, the Kimura 2 parameter value is replaced by the Ks value corresponding to the peaks.

### Pseudo gene analysis in S. tetrandra

For detecting pseudogenization events we first constructed blocks of synthenic genes between the *S. tetrandra* X and Y-chromosomes. Next, we searched for unpaired genes on both chromosomes in these blocks and searched for the corresponding protein against the region on the opposite chromosome (X to Y and vice versa) using exonerate [85]. Cases in which the matched protein overlapped with a predicted protein were discarded. For the remaining cases, pseudogenization events were identified as exonerate matches containing frameshifts or pre-mature termination codons.

## Declarations

### Ethics approval and consent to participate

Not applicable

### Consent for publication

Not applicable

### Availability of data and materials

Raw sequences and genome assemblies have been deposited and ENA database (https://www.ebi.ac.uk/ena/browser/home) under project number PRJEB59005.

### Authors’ contributions

YB and RF conceived the study. LK extracted plant DNA for ONT sequencing. RF and MPWK produced the genome assemblies. EIS, EvdW and MPWK performed the bioinformatic analyses. EIS, MPWK, CK, RT, RGFV and YB wrote the manuscript. All authors have read and accepted the final manuscript.

## Acknowledgements

We thank participating companies (Enza Zaden, Bejo Zaden B.V., Sakata, BASF and KWS) for providing plant materials and inputs to the research as well as the manuscript.

## Supplementary information

### Additional file 1 (word)

**AdditionalFile1.docx**

Supplementary Figures S1-S13

Supplementary Figures S16-S20

Supplementary Tables S1-S3

Supplementary Tables S11, S12

Appendix S1 and S2

### Additional file 2 (Excel workbook)

**AdditionalFile2.xlsx**

Supplementary Table S4-S10

### Additional file 3 (PDF)

**AdditionalFile3.pdf**

Supplementary Figure S14

### Additional file 4 (PDF)

**AdditionalFile4.pdf**

Supplementary Figure S15

