## Additional File 1 for "Chromosome level assembly of wild spinach provides insights into the divergence of homo- and heteromorphic plant sex-chromosomes"

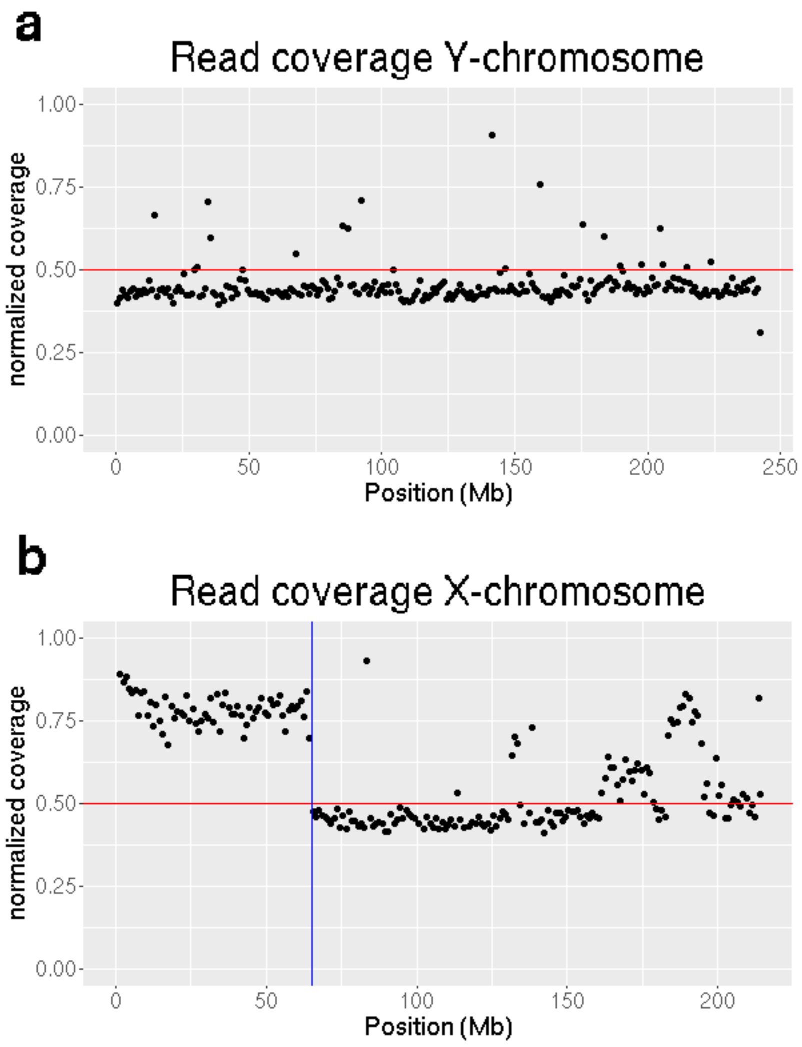


**Figure S1**. Illumina read coverage of a male *S. tetrandra* sample against the initial Y (a) and X (b) pseudo chromosomes. The red line corresponds to the expected diploid coverage across the chromosome. The blue vertical line corresponds to the region designated as the potential PSEUDO AUTOSOMAL REGION.

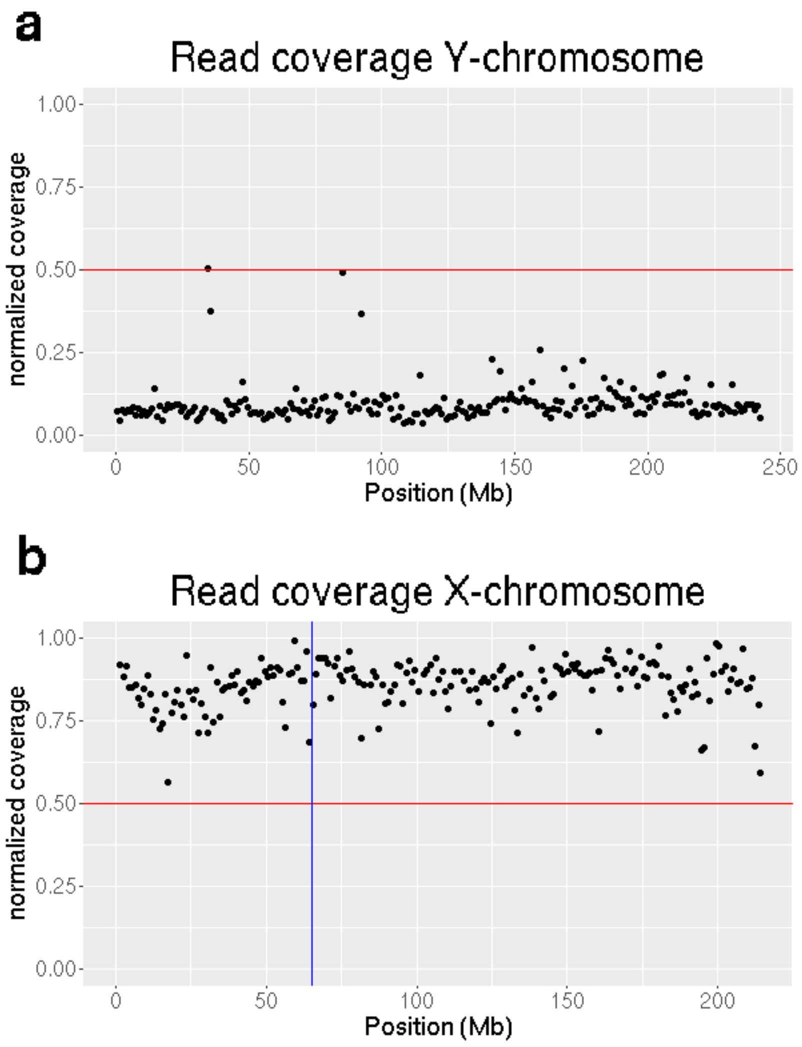


**Figure S2**. Illumina read coverage of a female *S. tetrandra* sample against the initial Y (a) and X (b) pseudo chromosomes. Horizontal red and vertical blue lines have the same interpretation as in Figure S1.

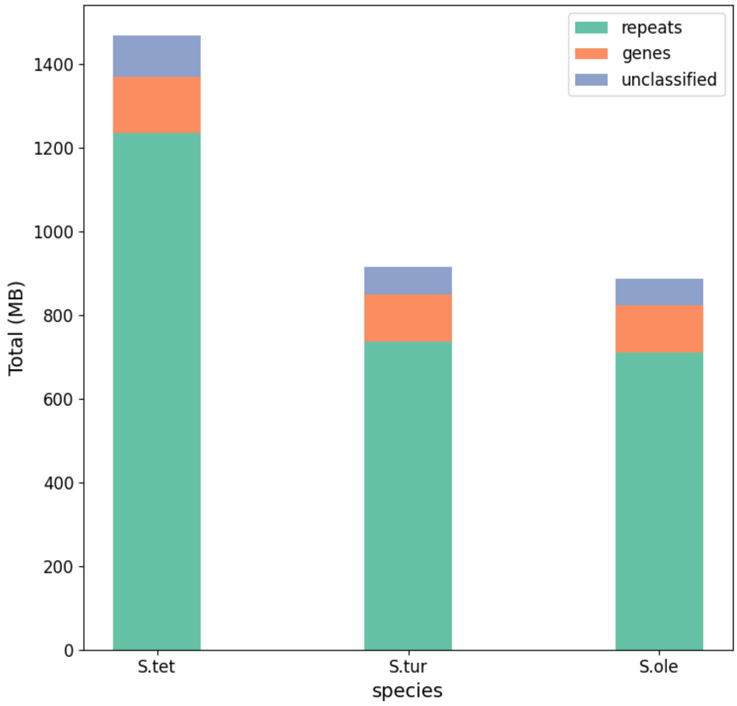


**Figure S3**. Total size of repeats and genes per genome. Length of individual genes were calculated from start to termination codon. Total repeat size corresponds to the number of bases masked by RepeatMasker. The remaining bases in the genome were labeled as unclassified.

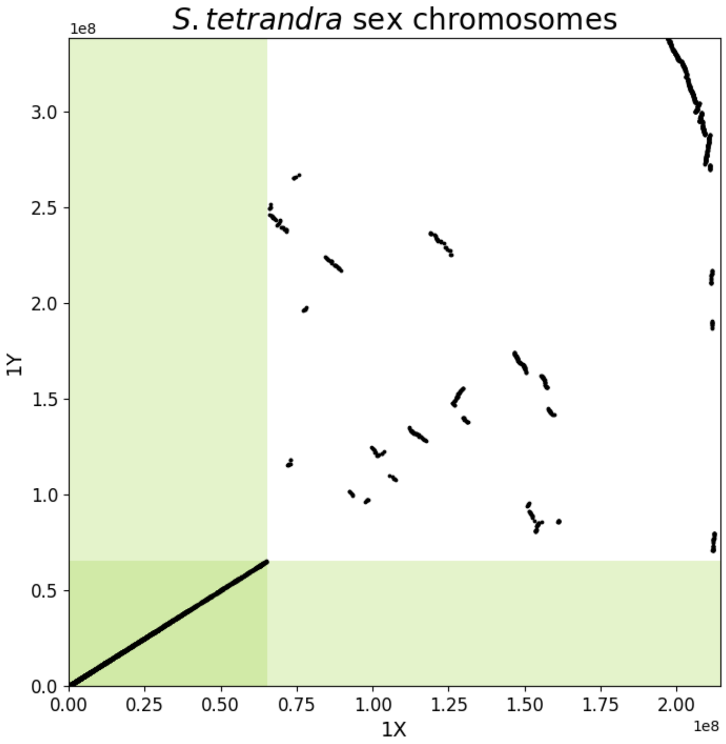


**Figure S4.** *S. tetrandra* sex chromosome comparison**.** Synthenic blocks between X- and Y-chromosomes were identified using MScanX. Green rectangles correspond to the predicted PSEUDO AUTOSOMAL REGIONS.

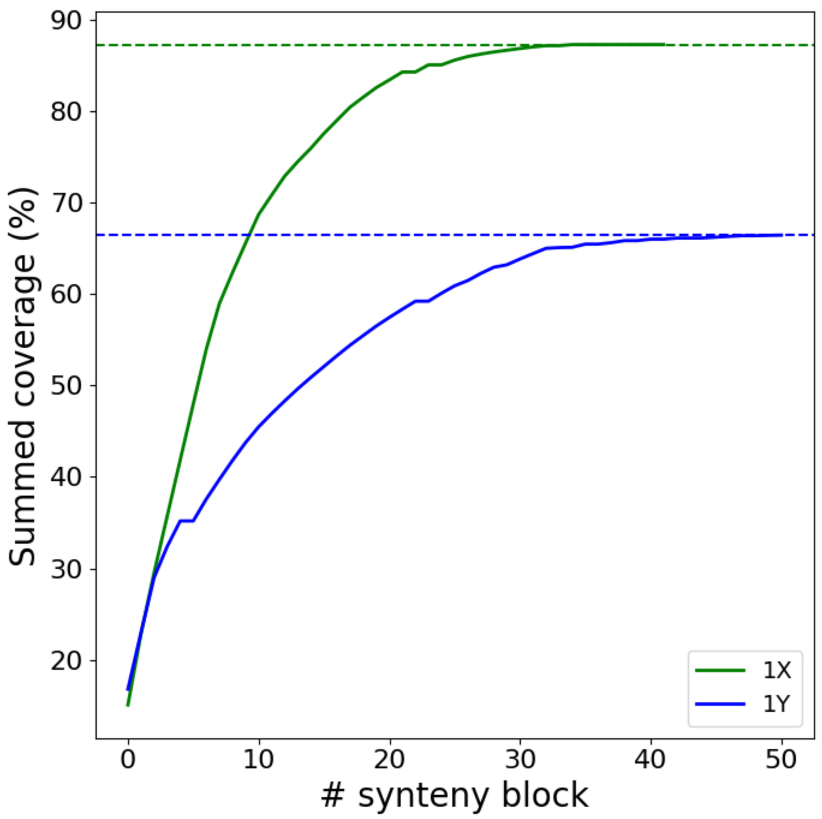


**Figure S5.** Fraction of *S. olereacea* chromosome 1X in syntenic blocks with *S. tetrandra* Sex chromosomes. Summed coverage (percentage) was obtained by first sorting the syntenic blocks from largest to smallest and then summing the percentage of the *S. oleracea* chromosome 1X covered by the blocks. Horizontal lines indicate the maximum percentage of the *S. oleracea* X chromosome that is covered by the syntenic blocks.

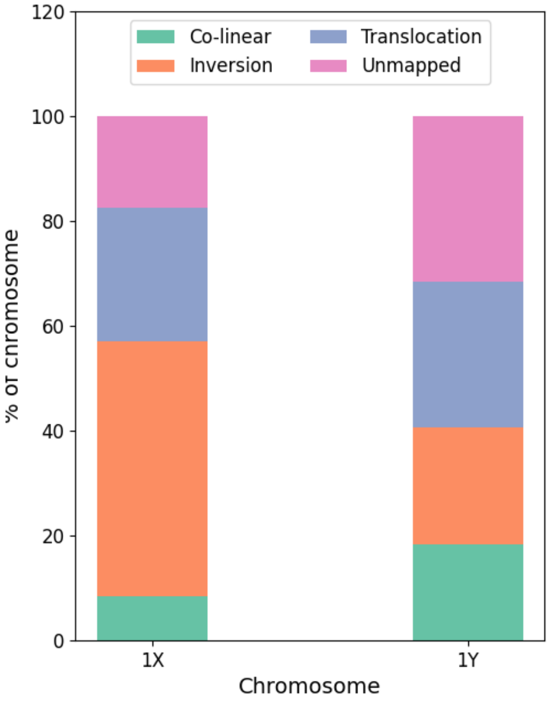


**Figure S6.** Rearrangement of *S. tetrandra* sex chromosomes relative to *S. oleracea* chromosome 1. Percentages represent the fraction of the X- or Y- chromosomes within a particular type of rearrangement.

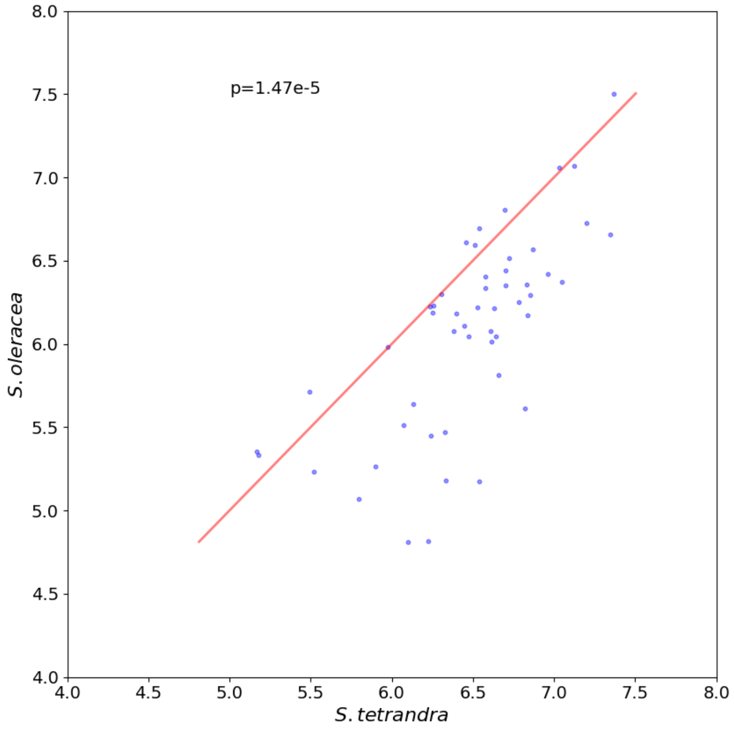


**Figure S7.** Size comparison of conserved segment sizes between *S. tetrandra* Y- and *S. oleracea* X chromosome. A sign-test (p-value in figure) was performed to determine whether the segments in one of the species is more often than expected larger.

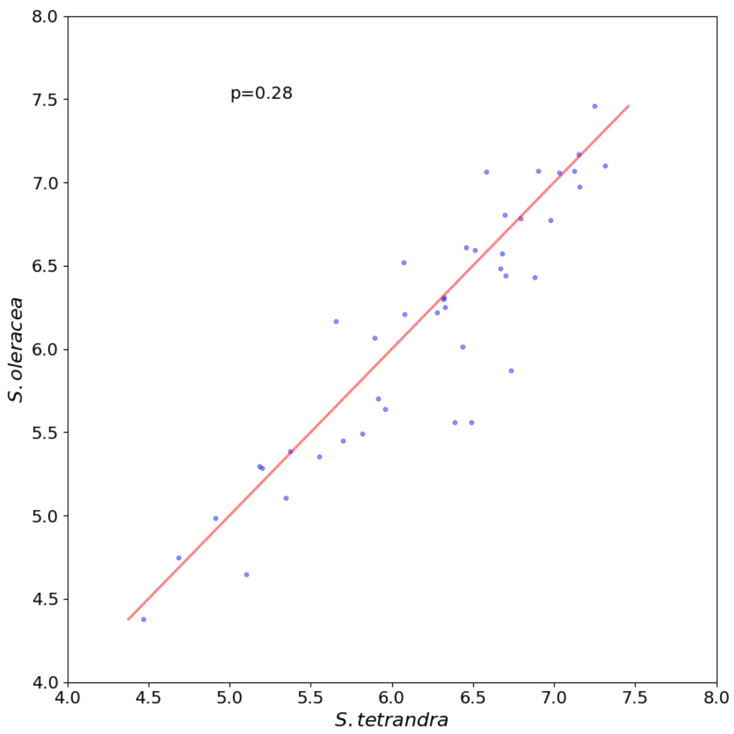


**Figure S8.** Size comparison of conserved segment sizes between *S. tetrandra* X- and *S. oleracea* X chromosome. A sign-test (p-value in figure) was performed to determine whether the segments in one of the species is more often than expected larger.

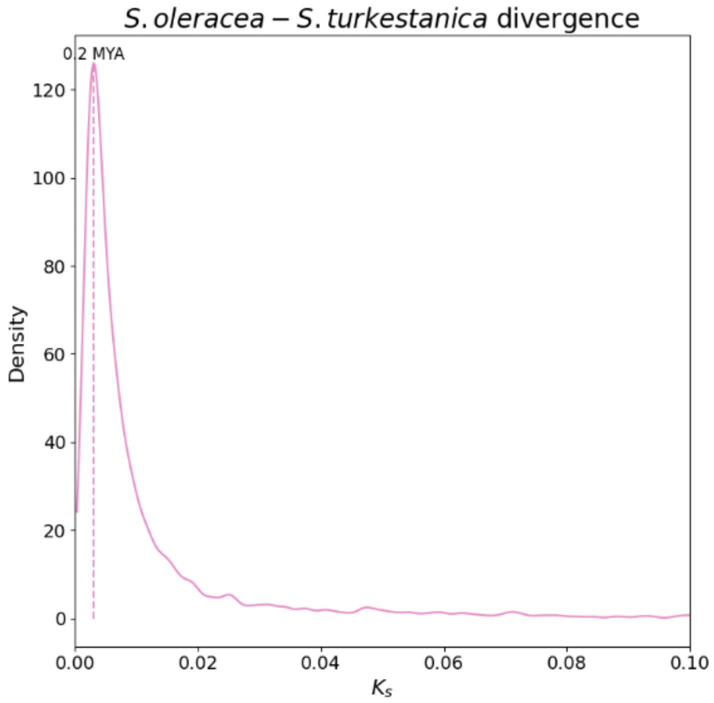


**Figure S9.** Divergence time between *S. turkestanica* and *S. oleracea* based on $K_s$ values**.** The divergence is derived from the mode of the $K_s$ distribution derived from autosomal ortholog gene pairs. A grid search was performed for finding the optimal bandwidth for the Gaussian kernel density fitting (Methods). Many $K_s$ values were exactly zero and were removed as they inhibited the identification of the peak.

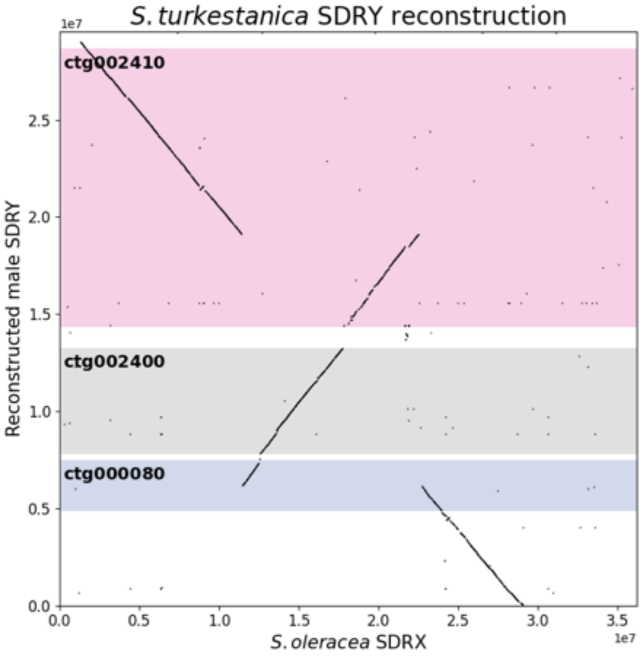


**Figure S10**. *S. turkestanica* SDR – Y assembly. The SDR region of *S. turkestanica* mainly consists of 3 larger contigs (shaded boxes). Contig000080 and Contig002410 capture inversion break points when compared to the female SDR X of *S. oleracea*.

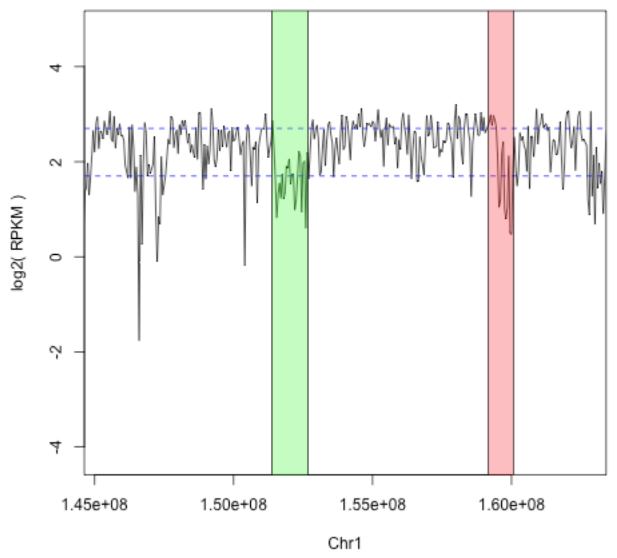


**Figure S11.** *S. turkestanica* nanopore coverage along the *S. oleracea* Y SDR. Top horizontal line corresponds to the expected diploid coverage measured as the mode from the log2 FPKM distribution values across bins on (50000nt) chromosome 1. The lower horizontal is obtained by subtracting 1 of the top line and corresponds to the expected haplotype coverage in log2 FPKM.

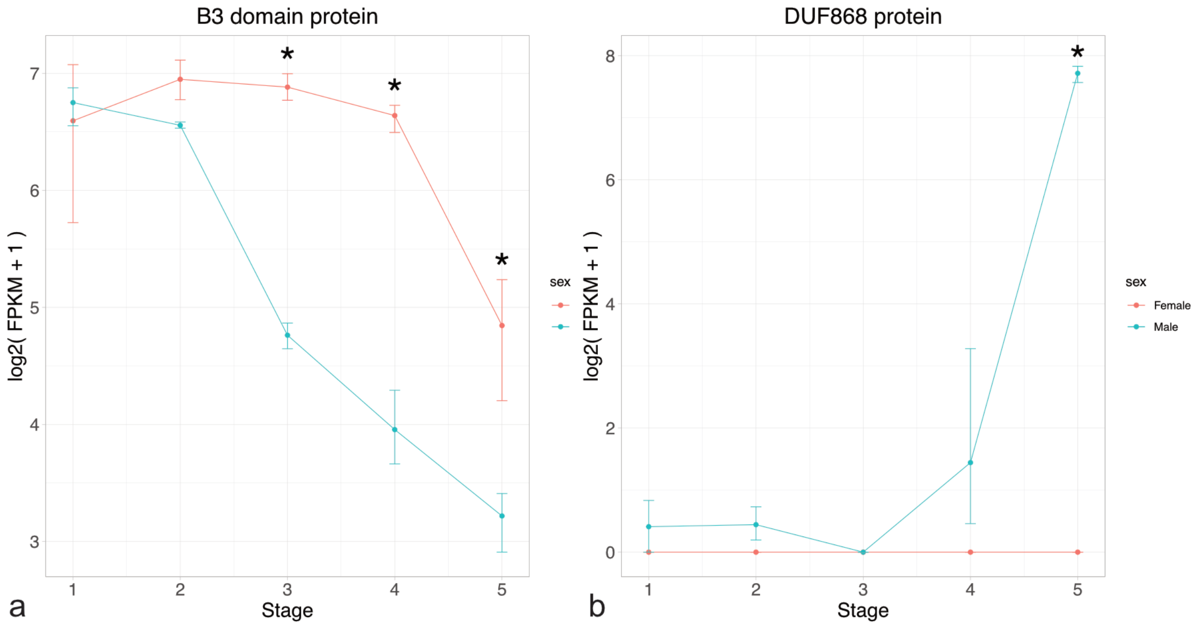


**Figure S12. *Expression of selected SDR genes during flower development in YY S. oleracea*.** **a.** Expression of the *S. oleracea YY22602* which a homolog *of the S. tetrandra* gene Stet|g4018 on chromosome X (Additional file 3: Fig S14). b. Expression of *S .oleracea YY22531* which is a homolog of *S. tetrandra Stet|g3474* on chromosome X (Additional file 4: Fig S15). Flower development stages correspond to those described in (13). Asterisks indicate the stages at which the expression level is significantly different between the male and female samples (adjusted p-value 0.01).

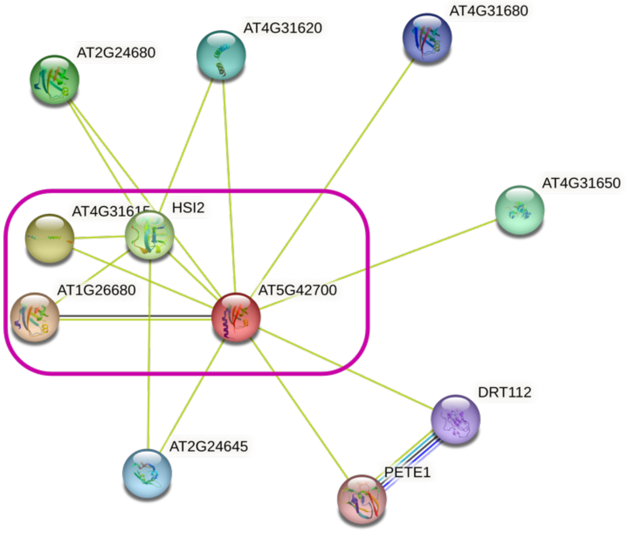


**Figure S13.** String based network around *AT5G42700*. Pink rectangle captures the single linkage cluster around *AT5G42700* between nodes linked with confidence values >0.7. The network is mostly derived by grouping genes that are more often than expected co-mentioned in PubMed abstracts.

### Figure S14 [Please refer to additional file 3]

### Figure S15 [Please refer to additional file 4]

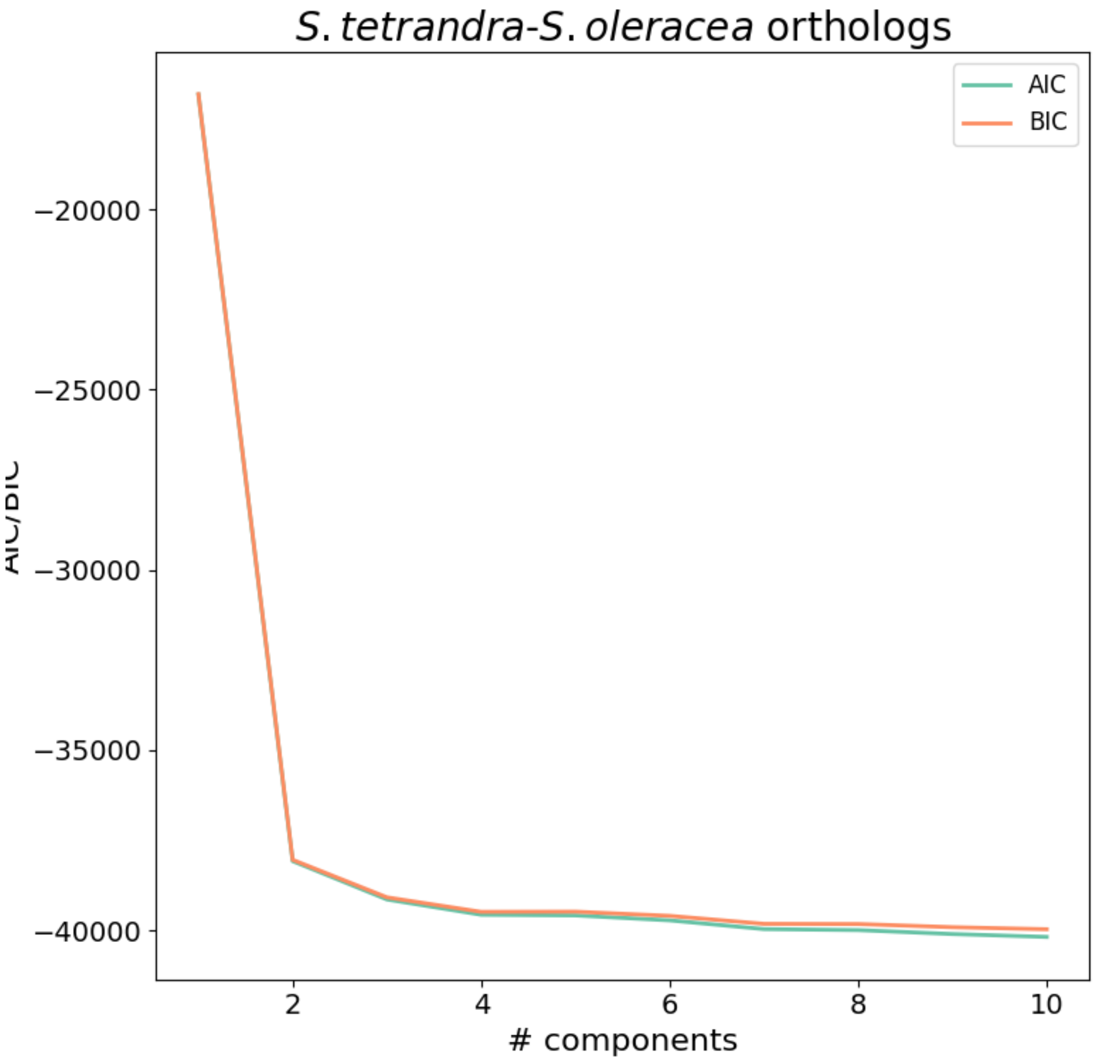


**Figure S16.** Selection of the number of Gaussian mixture model components in the $K_s$ distribution of *S. tetrandra – S. oleracea* autosomal ortholog pairs.

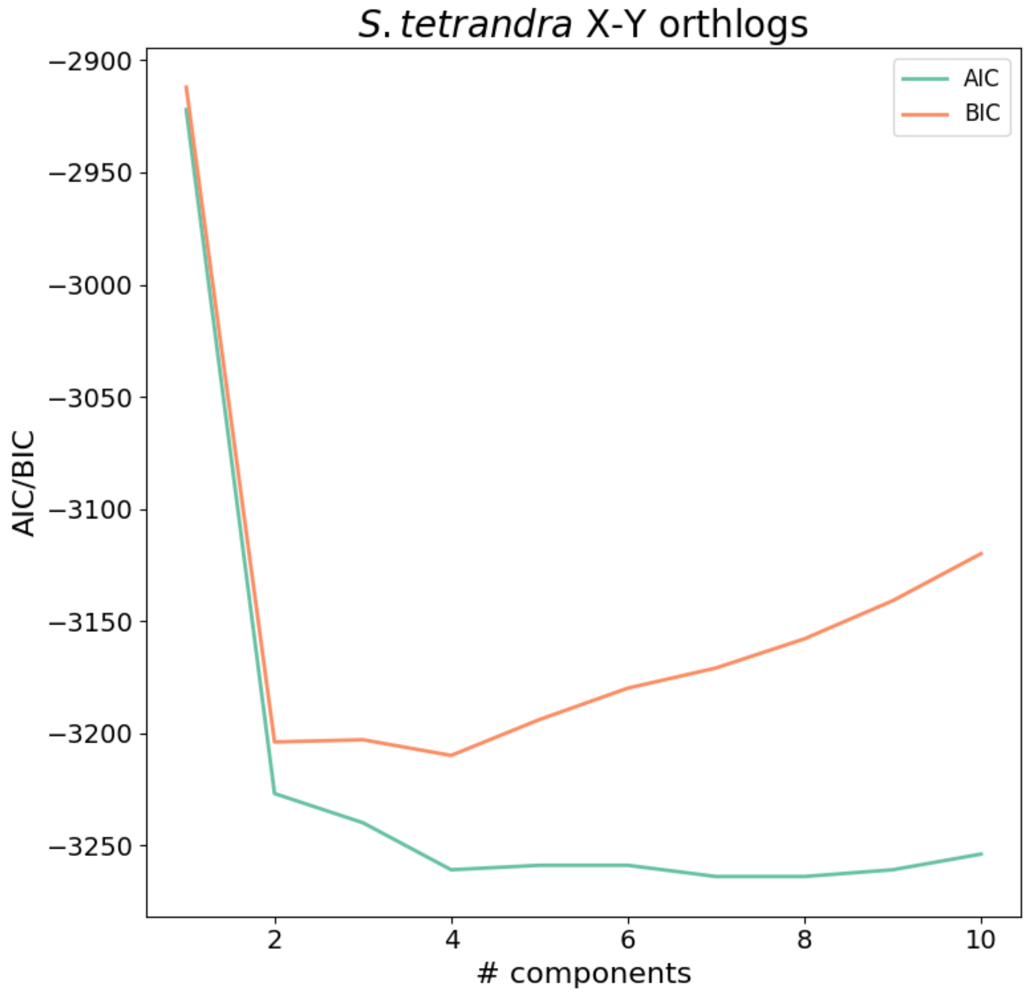


**Figure S17**. Selection of the number of Gaussian mixture model components underlying the $K_s$ distribution of *S. tetrandra* X – Y linked orthologs.

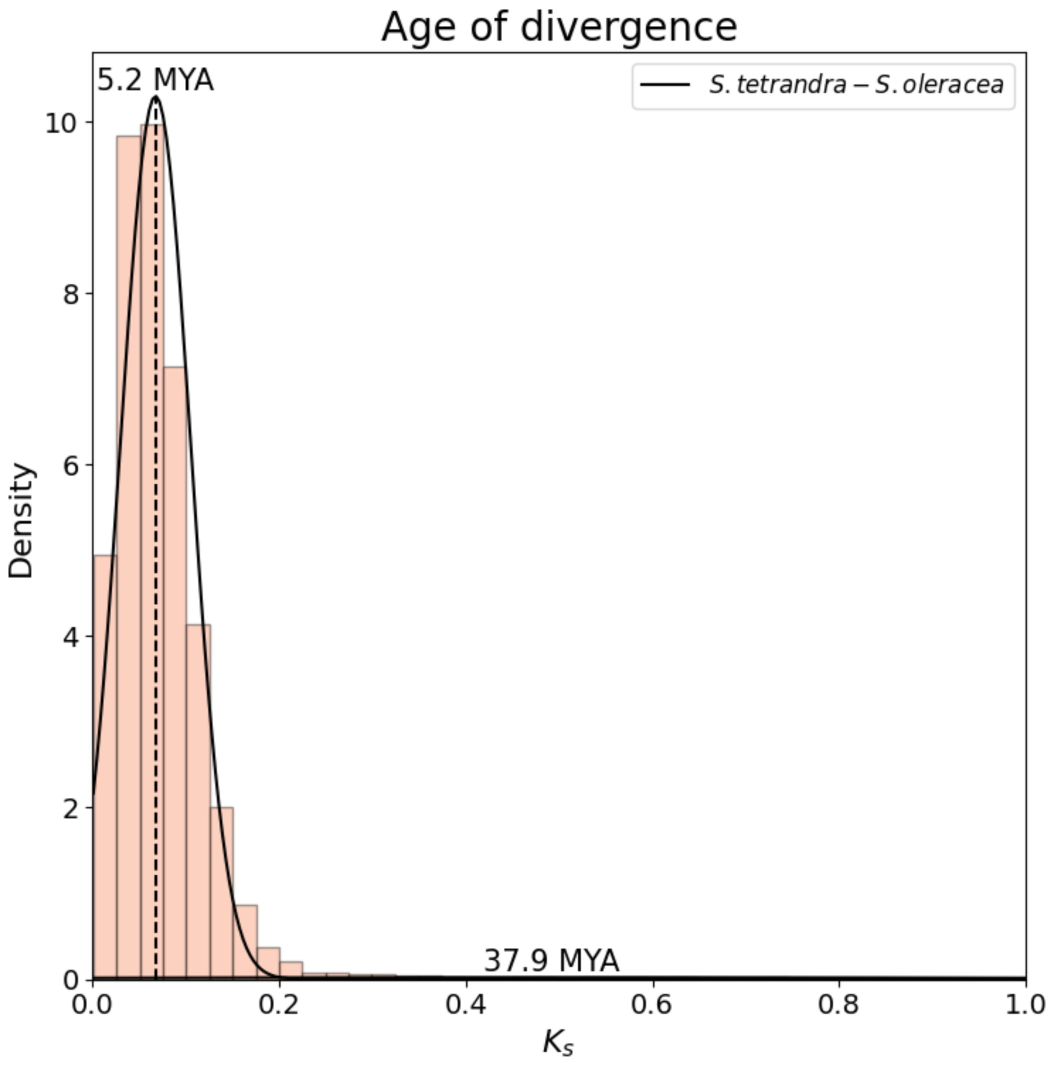


**Figure S18.** Fitting of Gaussian mixture model components onto the $K_s$ distribution of *S. tetrandra – S. oleracea* autosomal ortholog pairs. The empirical distribution is visualized as a histogram. Black lines correspond to the fitted components. The modes of the components were converted to age (see Methods section in the main text).

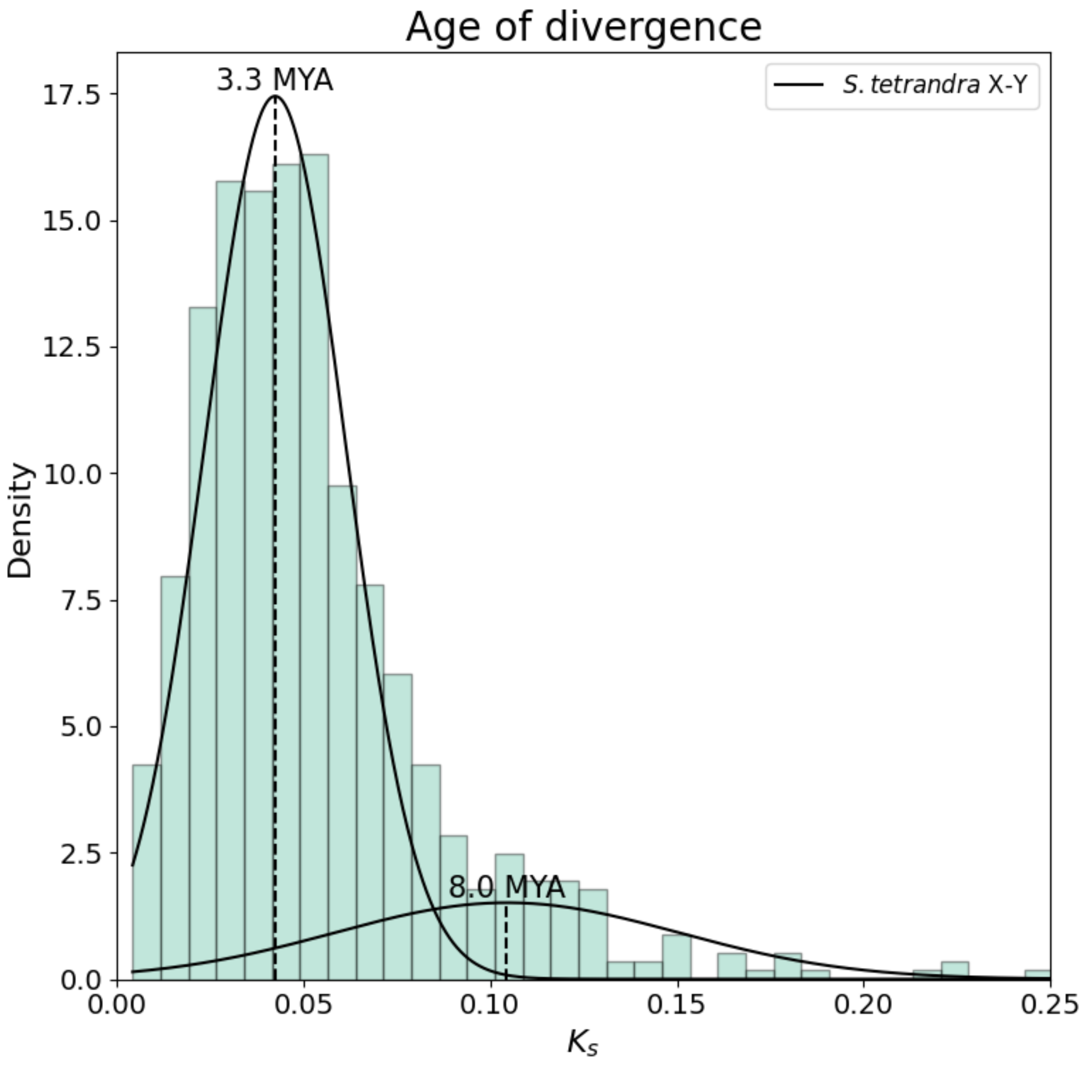


**Figure S19** Fitting of Gaussian mixture model components onto the $K_s$ distribution of *S. tetrandra* X-Y linked orthologs. The empirical distribution of $K_s$ values is displayed as a histogram on which the fitted models are plotted (black lines). The modes of the fitted Gaussian components were converted to age (see Methods section in main text).

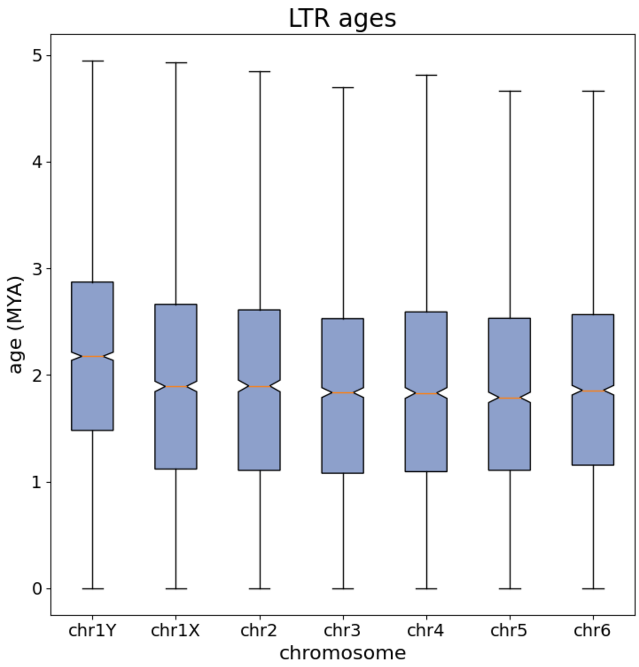


**Figure S20. Insertion age of *S. tetrandra* intact LTR per chromosome.** Notches correspond to the median of the distribution. Whiskers indicate the range of the data.

**Table S1.** Basic genome assembly and annotation properties.

| | *S. oleracea* | *S. turkestanica* | *S. tetrandra* |
|---|---|---|---|
| Estimated genome size (K-mer based) | 987072567 | 962791058 | 1198099674 |
| Total assembly size | 887785682 (89.9%) (894250018) | 917369818 (95.3%) | 1469268197 (122.6%) |
| Number of assembled pseudo chromosomes | 6 | 6 | 7 |
| Number of genes | 32029 | 33082 | 41548 |
| BUSCO score | 96.2% | 95.2% | 95.7% |
| Number of unique genes | 2396 (7.5%) | 2988 (9.0%) | 8530 (20.5%) |
| Repeat content (Mb & fraction) | 709591913 (79.9%) | 735766804 (80.2%) | 1235847581 (84.1%) |

**Table S2.** Full list of enriched (p <= 0.05) biological process GO terms for unique *S. tetrandra* genes located on the Y-chromosome. The terms were grouped using semantic similarity threshold of 0.45 (see Methods section in main text)

| GO-term | Description | Gene count | p-value |
|---|---|---|---|
| **GO:0006720** | **isoprenoid metabolic process (Top level)** | | |
| GO:0016103 | diterpenoid catabolic process | 12.0 | 0.00354695555555556 |
| GO:0045487 | gibberellin catabolic process | 12.0 | 0.00354695555555556 |
| GO:0016115 | terpenoid catabolic process | 12.0 | 0.00908366666666667 |
| GO:0008300 | isoprenoid catabolic process | 12.0 | 0.00908366666666667 |
| GO:0009685 | gibberellin metabolic process | 12.0 | 0.0263858888888889 |
| GO:0016101 | diterpenoid metabolic process | 12.0 | 0.0444914285714286 |
| **GO:0090304** | **nucleic acid metabolic process (Top level)** | | |
| GO:0071897 | DNA biosynthetic process | 33.0 | 1.187365e-11 |
| GO:0090501 | RNA phosphodiester bond hydrolysis | 31.0 | 8.5646e-09 |
| GO:0006259 | DNA metabolic process | 42.0 | 7.3967e-05 |
| GO:0090305 | nucleic acid phosphodiester bond hydrolysis | 38.0 | 0.012366 |
| **GO:0006382** | **adenosine to inosine editing (100%)** | | |
| GO:0002100 | tRNA wobble adenosine to inosine editing | 3.0 | 0.012366 |
| GO:0006382 | adenosine to inosine editing | 3.0 | 0.012366 |
| **Singletons** | | | |
| GO:0032197 | transposition, RNA-mediated | 30.0 | 1.84268666666667e-12 |
| GO:0032196 | transposition | 30.0 | 1.84268666666667e-12 |
| GO:0006508 | proteolysis | 43.0 | 0.00015572 |
| GO:0042447 | hormone catabolic process | 12.0 | 0.00716312 |
| GO:0022619 | generative cell differentiation | 3.0 | 0.012366 |
| GO:1903866 | palisade mesophyll development | 3.0 | 0.012366 |
| GO:0010377 | guard cell fate commitment | 3.0 | 0.0270461052631579 |
| GO:0072583 | clathrin-dependent endocytosis | 5.0 | 0.0291975 |

**Table S3**. Grouped enriched GO terms (p < 0.05) for genes originating through tandem duplications that are specific to *S. tetrandra*. Grouping of terms was done in the same way as for table S2.

| GO-term | Description | Gene count | p-value |
|---|---|---|---|
| **GO:0015855** | **pyrimidine nucleobase transport (57%)** | | |
| GO:0006863 | purine nucleobase transport | 10.0 | 2.10222e-06 |
| GO:1904823 | purine nucleobase transmembrane transport | 10.0 | 2.10222e-06 |
| GO:0042906 | xanthine transport | 4.0 | 0.000666135555555556 |
| GO:0015855 | pyrimidine nucleobase transport | 4.0 | 0.0290325423728814 |
| GO:0015857 | uracil transport | 4.0 | 0.0290325423728814 |
| GO:1903791 | uracil transmembrane transport | 4.0 | 0.0290325423728814 |
| GO:1904082 | pyrimidine nucleobase transmembrane transport | 4.0 | 0.0290325423728814 |
| **GO:0042221** | **response to chemical (Top level)** | | |
| GO:0046677 | response to antibiotic | 24.0 | 0.00295331034482759 |
| GO:0010045 | response to nickel cation | 4.0 | 0.00295331034482759 |
| GO:0010042 | response to manganese ion | 4.0 | 0.0105976111111111 |
| GO:0010043 | response to zinc ion | 8.0 | 0.0169013170731707 |
| GO:0070301 | cellular response to hydrogen peroxide | 6.0 | 0.0170205581395349 |
| GO:0042542 | response to hydrogen peroxide | 12.0 | 0.0170205581395349 |
| **GO:0015791** | **polyol transmembrane transport (80%)** | | |
| GO:0015795 | sorbitol transmembrane transport | 4.0 | 0.00164371111111111 |
| GO:0015797 | mannitol transmembrane transport | 4.0 | 0.00164371111111111 |
| GO:0015798 | myo-inositol transport | 4.0 | 0.00295331034482759 |
| GO:0015793 | glycerol transmembrane transport | 4.0 | 0.0060828125 |
| GO:0015791 | polyol transmembrane transport | 4.0 | 0.0167399 |
| **GO:0015750** | **pentose transmembrane transport (75%)** | | |
| GO:0015752 | D-ribose transmembrane transport | 4.0 | 0.00164371111111111 |
| GO:0015753 | D-xylose transmembrane transport | 4.0 | 0.00164371111111111 |
| GO:0015757 | galactose transmembrane transport | 4.0 | 0.00164371111111111 |
| GO:0015750 | pentose transmembrane transport | 4.0 | 0.00295331034482759 |
| **GO:0034471** | **ncRNA 5'-end processing (75%)** | | |
| GO:0034471 | ncRNA 5'-end processing | 6.0 | 0.0060828125 |
| GO:0000472 | endonucleolytic cleavage to generate mature 5'-end of SSU-rRNA from (SSU-rRNA, 5.8S rRNA, LSU-rRNA) | 4.0 | 0.0105976111111111 |
| GO:0000480 | endonucleolytic cleavage in 5'-ETS of tricistronic rRNA transcript (SSU-rRNA, 5.8S rRNA, LSU-rRNA) | 4.0 | 0.0105976111111111 |
| GO:0000967 | rRNA 5'-end processing | 4.0 | 0.0290325423728814 |
| **GO:0006857** | **oligopeptide transport (100%)** | | |
| GO:0042939 | tripeptide transport | 6.0 | 0.00113250909090909 |
| GO:0034635 | glutathione transport | 4.0 | 0.00295331034482759 |
| GO:0006857 | oligopeptide transport | 6.0 | 0.0119946486486486 |
| **GO:0006811** | **ion transport (50%)** | | |
| GO:0055085 | transmembrane transport | 59.0 | 0.00245259 |
| GO:0006811 | ion transport | 47.0 | 0.0182225531914894 |
| **GO:0019372** | **lipoxygenase pathway (100%)** | | |
| GO:0010597 | green leaf volatile biosynthetic process | 4.0 | 0.00295331034482759 |
| GO:0019372 | lipoxygenase pathway | 4.0 | 0.00295331034482759 |
| **GO:0006074** | **(1->3)-beta-D-glucan metabolic process (50%)** | | |
| GO:0051275 | beta-glucan catabolic process | 4.0 | 0.0218008 |
| GO:0006074 | (1->3)-beta-D-glucan metabolic process | 4.0 | 0.0383310769230769 |
| **GO:0090558** | **plant epidermis development (50%)** | | |

| | | | |
|---|---|---|---|
| GO:0090558 | plant epidermis development | 21.0 | 0.0254542307692308 |
| GO:0048468 | cell development | 28.0 | 0.0483675757575758 |
| **GO:0009804** | **coumarin metabolic process (100%)** | | |
| GO:0009804 | coumarin metabolic process | 4.0 | 0.0383310769230769 |
| GO:0009805 | coumarin biosynthetic process | 4.0 | 0.0383310769230769 |
| **Singletons** | | | |
| GO:0010143 | cutin biosynthetic process | 14.0 | 2.10222e-06 |
| GO:0015851 | nucleobase transport | 10.0 | 2.14115e-06 |
| GO:0072530 | purine-containing compound transmembrane transport | 10.0 | 9.03176e-06 |
| GO:0071366 | cellular response to indolebutyric acid stimulus | 6.0 | 2.3358e-05 |
| GO:0080026 | response to indolebutyric acid | 6.0 | 0.000300317142857143 |
| GO:0015720 | allantoin transport | 4.0 | 0.000666135555555556 |
| GO:0080027 | response to herbivore | 6.0 | 0.000747456 |
| GO:0010090 | trichome morphogenesis | 14.0 | 0.0011679 |
| GO:0043100 | pyrimidine nucleobase salvage | 4.0 | 0.00164371111111111 |
| GO:0051179 | localization | 106.0 | 0.00180307368421053 |
| GO:0009620 | response to fungus | 28.0 | 0.00295331034482759 |
| GO:0009395 | phospholipid catabolic process | 6.0 | 0.00295331034482759 |
| GO:0090626 | plant epidermis morphogenesis | 14.0 | 0.00441206666666667 |
| GO:0010026 | trichome differentiation | 14.0 | 0.00684224242424242 |
| GO:0050832 | defense response to fungus | 23.0 | 0.0120887894736842 |
| GO:0006995 | cellular response to nitrogen starvation | 7.0 | 0.015372358974359 |
| GO:0000053 | argininosuccinate metabolic process | 3.0 | 0.0176954545454545 |
| GO:0006855 | xenobiotic transmembrane transport | 14.0 | 0.0182225531914894 |
| GO:0042908 | xenobiotic transport | 14.0 | 0.0182225531914894 |
| GO:0006076 | (1->3)-beta-D-glucan catabolic process | 4.0 | 0.0218008 |
| GO:0019856 | pyrimidine nucleobase biosynthetic process | 4.0 | 0.0218008 |
| GO:0000966 | RNA 5'-end processing | 10.0 | 0.0254542307692308 |
| GO:0019375 | galactolipid biosynthetic process | 4.0 | 0.0290325423728814 |
| GO:0072337 | modified amino acid transport | 4.0 | 0.0290325423728814 |
| GO:0010207 | photosystem II assembly | 5.0 | 0.0383310769230769 |
| GO:0010421 | hydrogen peroxide-mediated programmed cell death | 4.0 | 0.0383310769230769 |
| GO:0072531 | pyrimidine-containing compound transmembrane transport | 4.0 | 0.0383310769230769 |

### Table S4-S10 [Please refer to additional file 2]

**Table S11**. Pairwise ks.tests with the alternative hypothesis that the cumulative distribution function of chromosome i (column) lies above that of chromosome j (row). P-values were corrected for multiple testing using the Benjamini-Hochberg method.

| | chr1X | chr1Y | chr2 | chr3 | chr4 | chr5 | chr6 |
|---|---|---|---|---|---|---|---|
| **chr1X** | - | 1.0 | 0.8179 | 0.0703 | 0.4363 | 0.1407 | 0.471 |
| **chr1Y** | 0.0 | - | 0.0 | 0.0 | 0.0 | 0.0 | 0.0 |
| **chr2** | 1.0 | 1.0 | - | 0.1783 | 0.34 | 0.0879 | 0.7191 |
| **chr3** | 1.0 | 1.0 | 1.0 | - | 1.0 | 0.891 | 1.0 |
| **chr4** | 1.0 | 1.0 | 1.0 | 1.0 | - | 0.8562 | 1.0 |
| **chr5** | 1.0 | 1.0 | 1.0 | 1.0 | 1.0 | - | 1.0 |
| **chr6** | 1.0 | 1.0 | 1.0 | 0.7277 | 1.0 | 0.471 | - |

**Table S12.** Dunn test results of LTR insertion ages per chromosome. Null hypothesis was rejected (green cells) at alpha (0.05) / 2. Multiple testing correction was done using the Benjamini-Hochberg method.

| | **chr1X** | **chr1Y** | **chr2** | **chr3** | **chr4** | **chr5** | **chr6** |
|---|---|---|---|---|---|---|---|
| **chr1X** | - | 0.0 | 0.3105 | 0.032 | 0.1591 | 0.0506 | 0.2422 |
| **chr1Y** | 0.0 | - | 0.0 | 0.0 | 0.0 | 0.0 | 0.0 |
| **chr2** | 0.3105 | 0.0 | - | 0.1661 | 0.324 | 0.1753 | 0.4219 |
| **chr3** | 0.032 | 0.0 | 0.1661 | - | 0.2974 | 0.4397 | 0.1821 |
| **chr4** | 0.1591 | 0.0 | 0.324 | 0.2974 | - | 0.3107 | 0.3517 |
| **chr5** | 0.0506 | 0.0 | 0.1753 | 0.4397 | 0.3107 | - | 0.2153 |
| **chr6** | 0.2422 | 0.0 | 0.4219 | 0.1821 | 0.3517 | 0.2153 | - |

**Appendix S1. Gene function description obtained from the STRING database in network around *AT3G19184.1.***

***AT3G19184.1***

AP2/B3-like transcriptional factor family protein; Its function is described as DNA binding; Involved in regulation of transcription, DNA-dependent; Located in cellular_component unknown; Expressed in 7 plant structures; Expressed during F mature embryo stage, petal differentiation and expansion stage, E expanded cotyledon stage, D bilateral stage; Contains the following InterPro domains: Transcriptional factor B3 (InterPro:IPR003340); BEST Arabidopsis thaliana protein match is: AP2/B3-like transcriptional factor family protein (TAIR:AT5G42700.1).

***AT3G17010.1, REM22***

Transcriptional factor B3 family protein, contains Pfam profile PF02362: B3 DNA binding domain. Activated by AGAMOUS ina a cal-1, ap1-1 background. Expressed in stamen primordia, the placental region of developing carpels and the ovary.

***AT1G26680.1***

Transcriptional factor B3 family protein; Its function is described as DNA binding, sequence-specific DNA binding transcription factor activity; Involved in regulation of transcription,

DNA-dependent; Located in chloroplast; Expressed in ovule, embryo; Expressed during D bilateral stage; Contains the following InterPro domains: Transcriptional factor B3 (InterPro:IPR003340); BEST Arabidopsis thaliana protein match is: Transcriptional factor B3 family protein (TAIR:AT2G24645.1).

***AT5G18000.1, VDD***

B3 domain-containing protein At5g18000; Encodes VERDANDI (VDD), a putative transcription factor belonging to the reproductive meristem (REM) family. VDD is a direct target of the MADS domain ovule identity complex. Mutation in VDD affects embryo sac differentiation.

***AT3G53310.1***

AP2/B3-like transcriptional factor family protein; Its function is described as DNA binding, sequence-specific DNA binding transcription factor activity; Involved in regulation of transcription, DNA-dependent; Located in endomembrane system; Expressed in 15 plant structures; Expressed during 13 growth stages; Contains the following InterPro domains: Transcriptional factor B3 (InterPro:IPR003340).

**Appendix S2. Gene function description obtained from the STRING database in network around *AT5G42700***

***AT5G42700***

AP2/B3-like transcriptional factor family protein; Its function is described as DNA binding; Involved in regulation of transcription, DNA-dependent; Located in cellular_component unknown; Expressed in 8 plant structures; Expressed during 4 anthesis, F mature embryo stage, petal differentiation and expansion stage, E expanded cotyledon stage, D bilateral stage; Contains the following InterPro domains: Transcriptional factor B3 (InterPro:IPR003340); BEST Arabidopsis thaliana protein match is: AP2/B3-like transcriptional factor family protein (TAIR:AT3G19184.1).

***AT1G26680***

Transcriptional factor B3 family protein; Its function is described as DNA binding, sequence-specific DNA binding transcription factor activity; Involved in regulation of transcription, DNA-dependent; Located in chloroplast; Expressed in ovule, embryo; Expressed during D bilateral stage; Contains the following InterPro domains: Transcriptional factor B3 (InterPro:IPR003340); BEST Arabidopsis thaliana protein match is: Transcriptional factor B3 family protein (TAIR:AT2G24645.1).

***AT4G31615***

Transcriptional factor B3 family protein; Its function is described as DNA binding, sequence-specific DNA binding transcription factor activity; Involved in regulation of transcription, DNA-dependent; Located in cellular_component unknown; Expressed in shoot apex, embryo, leaf whorl, flower, seed; Expressed during F mature embryo stage, petal differentiation and expansion stage, E expanded cotyledon stage, D bilateral stage; Contains the following InterPro domains: Transcriptional factor B3 (InterPro:IPR003340).

***HSI2, AT2G30470***

B3 domain-containing transcription repressor VAL1; HSI2 is a member of a novel family of B3 domain proteins with a sequence similar to the ERF-associated amphiphilic repression (EAR) motif. It functions as an active repressor of the Spo minimal promoter (derived from a gene for sweet potato sporamin A1) through the EAR motif. It contains a plant-specific B3 DNA-binding domain. The Arabidopsis genome contains 42 genes with B3 domains which could be classified into three families that are represented by ABI3, ARF1 and RAV1
