## Additional File 3 for "Chromosome level assembly of wild spinach provides insights into the divergence of homo- and heteromorphic plant sex-chromosomes"

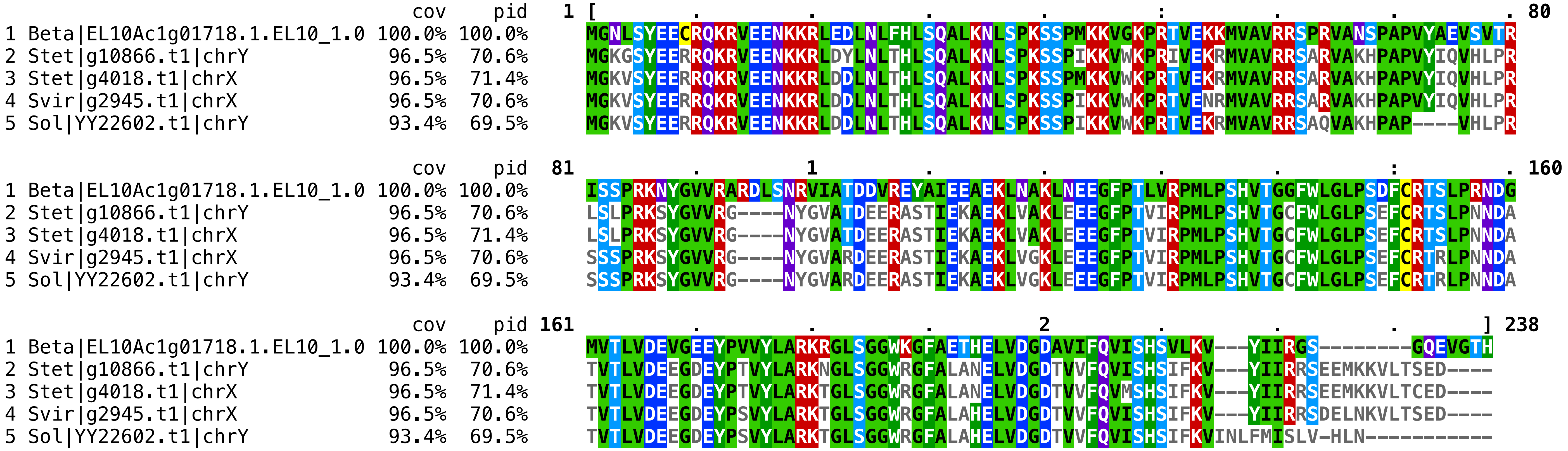


**Figure S14. Protein alignment of B3 domain containing proteins.** (Beta) *B. vulgaris*,(Stet) *S. tetrandra*. (Svir). *S. oleracea viroflay*, (Sol) *S. oleracea YY*
