## Additional File 4 for "Chromosome level assembly of wild spinach provides insights into the divergence of homo- and heteromorphic plant sex-chromosomes"

```
                                    cov    pid   1 [        .         .         .         .         :         .         .         . 80
1 Beta|EL10Ac9g20664.1.EL10_1.0 100.0% 100.0%     ---------MKGPPERITEEPMTLKSSLCTVSSVFLTNIAGYWRNVSVLWCKNMMNISFSIIIDGFEGDLETQTTCKVEL
2 Stet|g12725.t1|chrY             99.7%  82.4%     MTTAIRARSLRGPPERITEEPITLKSSLCTVTSVFLTNIAGYWRNVSVLWCKNLMNISFSIIIDGFEGEMERQSTCKVEL
3 Svir|g3123.t1|chrX              99.7%  82.4%     MTTAIRARSLRGPPERITEEPITLKSSLCTVTSVFLTNIAGYWRNVSVLWCKNLMNISFSIIIDGFEGEMERQSTCKVEL
4 Sol|YY22531.t1|chrY             99.7%  82.4%     MTTAIRARSLRGPPERITEEPITLKSSLCTVTSVFLTNIAGYWRNVSVLWCKNLMNISFSIIIDGFEGEMERQSTCKVEL
5 Stet|g3474.t1|chrX              99.7%  81.8%     MTSGIRAGSLRGPPERITEEPITLKSSLCTVTSVFLTNIAGYWRNVSVLWCKNLMNISFSIIIDGFEGEMERQSICKVEL

                                    cov    pid  81          .         1         .         .         .         .         :         . 160
1 Beta|EL10Ac9g20664.1.EL10_1.0 100.0% 100.0%     KPWHFWSRKGYKSFEINIQHKLAEESYHQLDIYWDLRSAKFSGSSPEPCSDYYLALVSDEEVVLLLGDNKKKAYKRTKKR
2 Stet|g12725.t1|chrY             99.7%  82.4%     KPWHFWSKKGYKTFEMD-QNKLAEEAGHQLDVYWDLRSAKFAGSSPEPCSDYYVALVSDEEVVLLLGDFKKKAYKRTKKR
3 Svir|g3123.t1|chrX              99.7%  82.4%     KPWHFWSKKGYKSFEMD-QNKLAEEAGHQLDVYWDLRSAKFAGSSPEPCSDYYVALVSDEEVVLLLGDFKKKAYKRTKKR
4 Sol|YY22531.t1|chrY             99.7%  82.4%     KPWHFWSKKGYKSFEMD-QNKLAEEAGHQLDVYWDLRSAKFAGSSPEPCSDYYVALVSDEEVVLLLGDFKKKAYKRTKKR
5 Stet|g3474.t1|chrX              99.7%  81.8%     KPWHFWSKKGYKSFEMD-QNKLAEEAGHQLDVYWDLRSAKFAGSSPEPCSDYYVALVSDEEVVLLLGDFKKKAYKRTKKR

                                    cov    pid 161          .         .         .         2         .         .         .         . 240
1 Beta|EL10Ac9g20664.1.EL10_1.0 100.0% 100.0%     PALVDAIQSLKKEIVYAKKSFATRARFTEGKGEHDVVIESSTSGPKDPEMWISIDGIVLIHVKNLQWKFRGNEMVVVNKT
2 Stet|g12725.t1|chrY             99.7%  82.4%     PALVDAIQSLKKETVYAKKSFSTRARLTENKWEHDVVVESSTTGSKDPEMWISIDGIVLIHIKNLQWKFRGNETVVVNKT
3 Svir|g3123.t1|chrX              99.7%  82.4%     PALVDAIQSLKKETVYAKKSFSTRARLTENKWEHDVVVESSTTGSKDPEMWISIDGIVLIHIKNLQWKFRGNETVVVNKT
4 Sol|YY22531.t1|chrY             99.7%  82.4%     PALVDAIQSLKKETVYAKKSFSTRARLTENKWEHDVVVESSTTGSKDPEMWISIDGIVLIHIKNLQWKFRGNETVVVNKT
5 Stet|g3474.t1|chrX              99.7%  81.8%     PALVDAIQSLKKETVYAKKSFSTRARLTENKWEHDVVVESSTTGSKDPEMWISIDGIVLIHIKNLQWKFRGNETVVVNKT

                                    cov    pid 241          :         .         .         .         .         3         .       ] 319
1 Beta|EL10Ac9g20664.1.EL10_1.0 100.0% 100.0%     PIQVLWDVHAWLFSNPGTGCGLFVFKPGAPEVEFDREG---SNRSGNSDNSFGSMFYSTQSTQTKIVDFCLLLQAWKID
2 Stet|g12725.t1|chrY             99.7%  82.4%     PVQVLWDVHAWLFCSPGTGYGLFIFKPGTPEVDTDKEGSKSSNRSGASDNSCASMFYSTQSTQTKIVDFCLLLQAWKTD
3 Svir|g3123.t1|chrX              99.7%  82.4%     PVQVLWDVHAWLFCSPGTGYGLFIFKPGTPEVDTDKEGSKSSNRSGASDNSCASMFYSTQTTQTKIVDFCLLLQAWKTD
4 Sol|YY22531.t1|chrY             99.7%  82.4%     PVQVLWDVHAWLFCSPGTGYGLFIFKPGTPEVDTDKEGSKSSNRSGASDNSCASMFYSTQTTQTKIVDFCLLLQAWKTD
5 Stet|g3474.t1|chrX              99.7%  81.8%     PVQVLWDVHAWLFCSPGTGYGLFIFKPGTPEVDTDKEGSKSSNRSGASDNTCASMFYSTQTTQTKIVDFCLLLQAWKTD
```

**Figure S15. Alignment of DUF868 domain containing proteins.** (Beta) *B. vulgaris* ,(Stet) *S. tetrandra*. (Svir). *S. oleracea viroflay*, (Sol) *S. oleracea YY*
